# Hippocampal mechanisms underlying the resolution of competition in memory and perception

**DOI:** 10.1101/2023.10.09.561548

**Authors:** Serra E. Favila, Mariam Aly

**Author notes:** Please address correspondence to: Serra Favila.

## Abstract

Behaving adaptively requires selection of relevant memories and sensations and suppression of competing ones. We hypothesize that these mechanisms are linked, such that hippocampal computations that resolve competition in memory also shape the precision of sensory representations to guide selective attention. We leverage fMRI-based pattern similarity, receptive field modeling, and eye tracking to test this hypothesis in humans performing a memory-dependent visual search task. In the hippocampus, more distinct representations of competing memories are associated with the accuracy of memory-guided eye movements. In visual cortex, preparatory coding of remembered target locations precedes search successes, whereas preparatory coding of competing locations precedes search failures due to interference. These effects are linked: more distinct hippocampal memories are associated with lower competitor activation in visual cortex, yielding more precise preparatory representations. These results demonstrate a role for memory distinctiveness in shaping the precision of sensory representations, highlighting links between mechanisms that overcome competition in memory and perception.

## Introduction

At any given moment, we are faced with multiple sensations, memories, and thoughts that compete to take control of our behavior. Behaving adaptively requires resolution of this competition, to focus on only those memories or external events that are relevant in the current context. Multiple cognitive domains, from perception to memory, have highlighted the necessity of selecting between competing representations for generating adaptive behavior. Yet, these aspects of cognition are largely studied separately, leading to assumptions that the way competition is resolved in memory is fundamentally different from how it is resolved in perception. An alternative possibility is that the computational principles that overcome competition in memory and perception are closely intertwined, such that mechanisms that resolve competition in memory also shape the precision of online perceptual behavior.

Within the field of visual perception, it has long been appreciated that limited processing capacity requires humans to make moment-by-moment decisions about what information to attend to and what to ignore^1,2^. These competitive interactions have been well-characterized in behavior and within visual cortex, where attention is known to modulate the strength and precision of stimulus representations^3–8^. Separately, within the field of episodic memory, a considerable amount of research has focused on the role of the hippocampus in differentiating similar memories such that competition between them is minimized. Both computational models^9,10^ and numerous fMRI studies in humans^11–16^ have shown that the hippocampus can learn to represent competing memories more dissimilarly than non-competing memories, with this memory differentiation serving to reduce interference in behavior. Although competitive interactions in attentional selection and memory are typically studied independently and in separate parts of the brain, there is reason to expect that these mechanisms are linked via the hippocampus. Prior work has shown that the hippocampus is involved in online attentional behavior^17–22^, codes for gaze-related variables at the single cell level^23–25^, and interacts with visual and oculomotor systems in support of memory-guided attention^26–31^. Combining these disparate literatures, we hypothesized that hippocampal mechanisms that make competing memories distinct would be associated with precise attentional selection and reduced activation of competing features in visual cortex.

In this work, we introduce a task that requires participants to rely on competing memories to guide visual attention. Participants first acquire pairs of competing memories, which consist of two highly similar scenes that are associated with distinct spatial locations. Then, while being scanned with fMRI, participants perform a difficult visual search task. Critically, the search task is structured such that participants can use memories from the previous session to predict the location of upcoming search targets. We record participants’ eye movements continuously during this task, allowing us to assess the efficiency of attentional guidance. Using a combination of pattern similarity and population receptive field modeling approaches to analyze the fMRI data, we measure: 1) the separability of competing memories in the hippocampus; and 2) preparatory representations of target and competitor locations in visual cortex prior to search onset. By combining these fMRI measures with our eye-tracking based behavioral measurements, we link variability in hippocampal separability of competing memories to the precision of visual cortical representations and to the accuracy of memory-guided eye movements during visual search.

## Results

### Competitive scene-location learning

Human participants (N = 32) took part in a two-session experiment that spanned consecutive days. In session 1, participants learned 24 scene-location associations. To create memory competition, the 24 scenes consisted of 12 pairs of highly similar images (pairmates; see Fig. 1A for examples). Each scene was associated with a spatial location, which we will refer to as its target location (Fig. 1B). There were 8 possible locations arranged in a circle, equidistant from the center of the screen and equidistant from each other. Pairmates were never associated with the same location. Participants were first exposed to the scenes without their associated locations and then performed interleaved study and test blocks on the scene-location associations (Fig. 1C). During study blocks, participants were presented with the scene-location associations, one at a time. During test blocks, participants were cued with one of the scene images and asked to respond by making a memory-guided saccade to the associated target location on a blank display. To succeed at this task, participants had to discriminate between scene pairmates such that they could respond with the target location associated with the cued scene and avoid responding with the competitor location associated with the cued scene’s pairmate.

**Figure 1.**
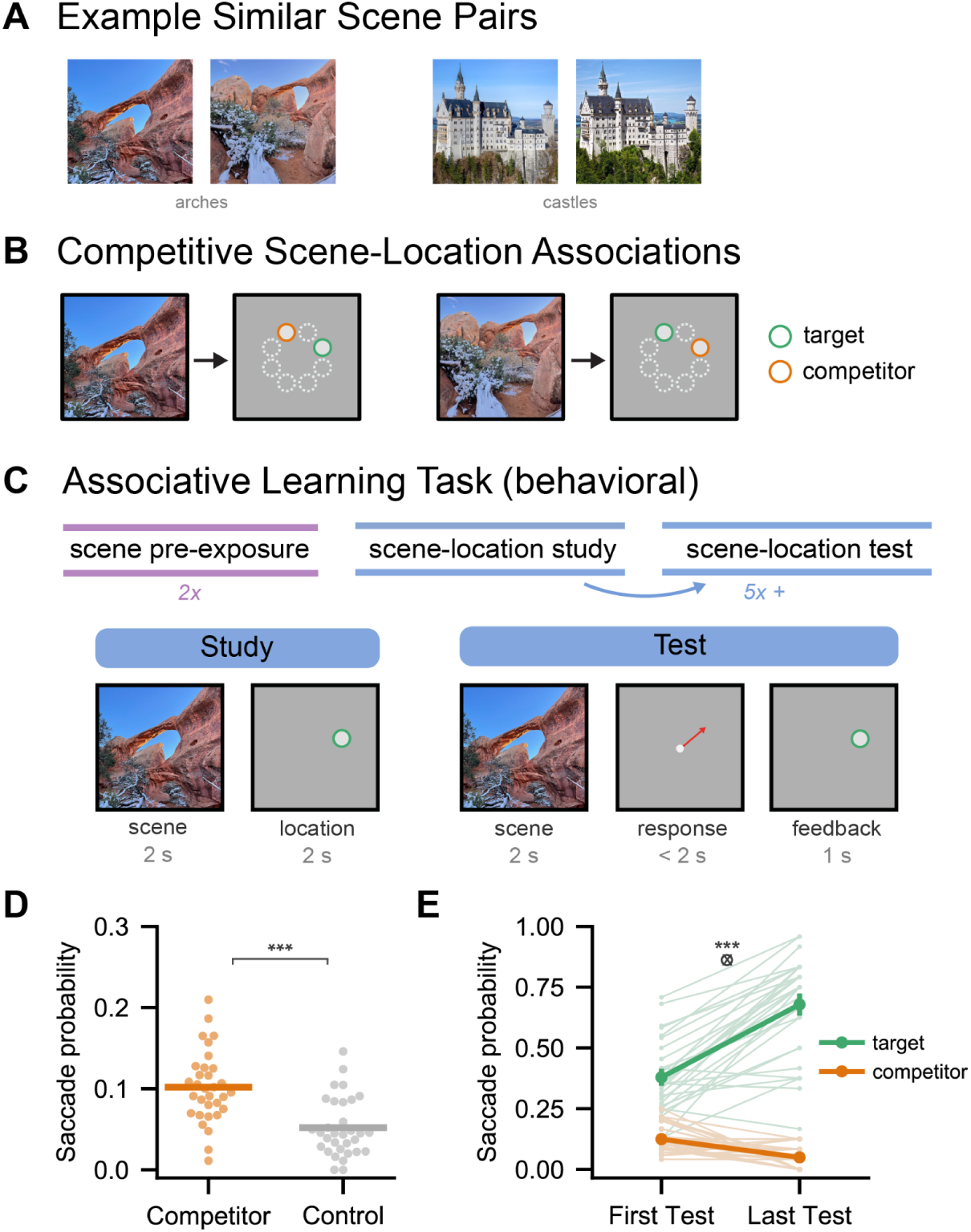
Experimental design and behavioral data from competitive learning task (session 1). **(A)** Representative example pairs of highly similar scenes (pairmates) comparable to those used in the experiment. **(B)** Each scene was associated with a target location, which was one of eight possible spatial locations. The pairmate structure of the scenes was intended to create memory competition, such that participants would bring to mind the competitor location (the target location of the similar scene). The competitor location is highlighted for visualization; participants saw each image paired exclusively with its target location. **(C)** Participants (N = 32) were exposed to all 24 scenes two times each. They then acquired the scene-location associations through interleaved study and test blocks. During study trials, participants intentionally encoded the scene-location pairs. During test trials, participants were cued with a scene and after a brief central fixation interval (not shown), made a memory-guided saccade to the recalled location. Participants were shown the correct location at the end of the trial as feedback. **(D)** Participants were more likely to make saccades to the competitor location than to a control location equidistant from the target (t(31) = 4.74, p < 0.001, d_z_ = 1.25, competitor - control = 0.05, 95% CI = [0.03, 0.07]; N = 32 participants; two-tailed paired t-test), establishing the presence of memory competition. Dots represent individual participants and lines represent the across-participant mean. **(E)** Participants’ memory improved over the course of the session, reflected by more frequent saccades to the target location and less frequent saccades to the competitor location (F(1,31) = 95.8, p < 0.001, η^2^_P_ = 0.76; N = 32 participants; repeated measures ANOVA). Light lines represent individual participants. Dark lines represent the across-participant mean ± SEM. *** p < 0.001.

To evaluate memory, we measured the angular error between participants’ final saccade end point and three relevant locations along the circle of studied locations: 1) the target location; 2) the competitor location; and 3) a control location an equal distance away from the target as the competitor. It was not possible to define a control location when target-competitor distances were 180 degrees, and thus these trials were excluded for all control vs. competitor location comparisons. Note that for all analyses, the term degrees indicates the polar angle distance, or distance around the unit circle, and not degrees of visual angle. We assigned a participant’s final saccade to one of these three locations if it was closer to that location than to any of the other studied locations (i.e. if the angular error between the saccade end point and the center of the relevant location was less than 22.5 degrees). To establish whether participants experienced memory competition, we first examined the relative probability of saccades to the competitor and control locations. Across all test trials, participants were significantly more likely to make responses to the competitor than to the control location (t(31) = 4.74, p < 0.001, d_z_ = 1.25, competitor - control = 0.05, 95% CI = [0.03, 0.07]; Fig. 1D), validating the presence of competition. We then confirmed that learning occurred during the session by comparing target and competitor response probabilities between the first and last test presentation. The relative probability of these responses changed reliably over the session (target/competitor x first/last interaction: F(1,31) = 95.8, p < 0.001, η^2^_P_ = 0.76; Fig. 1E). Participants made significantly more responses to the target location from the first to the last test presentation (t(31) = 9.82, p < 0.001; d_z_ = 1.74, last – first = 0.30%, 95% CI = [0.24, 0.36]) and significantly fewer responses to the competitor (t(31) = −5.60, p < 0.001, d_z_ = 1.49, last - first = −0.075%, 95% CI = [−0.10, −0.05]). Thus, participants successfully acquired the scene-location associations in the first experimental session, overcoming interference from memory competitors to do so.

### Memory-guided visual search behavior

In session 2, participants performed a memory-guided visual search task while being scanned with fMRI. On every trial, participants were briefly presented with a scene image (1.25 s) that contained a distortion (Fig. 2A; see Methods/Scene Distortions). Participants’ task was to find this distortion as quickly and accurately as possible by moving their eyes and holding fixation on the distortion once they found it. Critically, we embedded structure in the task that allowed participants to use the memories they formed in session 1 to boost their performance. Participants were told that the location associated with a given scene from session 1 was a reliable predictor of the distortion location on the next trial. A long intertrial interval (ITI) and brief scene presentations encouraged participants to engage in memory-based prediction to allocate attention optimally on the next trial. Participants were asked to fixate centrally during the ITI so that we could measure preparatory neural representations in retinotopic cortex. To test whether participants were actually engaging in a memory-based strategy, we manipulated the validity of memory-based prediction, following a well-established attentional cueing paradigm^32^. There were three critical task conditions (Fig. 2B). On valid trials (75% of trials), the distortion was placed in the predicted target location. On invalid trials (12.5% of trials), the distortion was placed in the competitor location. On no-prediction trials (12.5% of trials), the distortion was placed randomly in one of the 8 possible locations used in the experiment (Fig. 1B). These no-prediction trials always followed trial-unique novel scenes that had no associated location that participants could recall.

**Figure 2.**
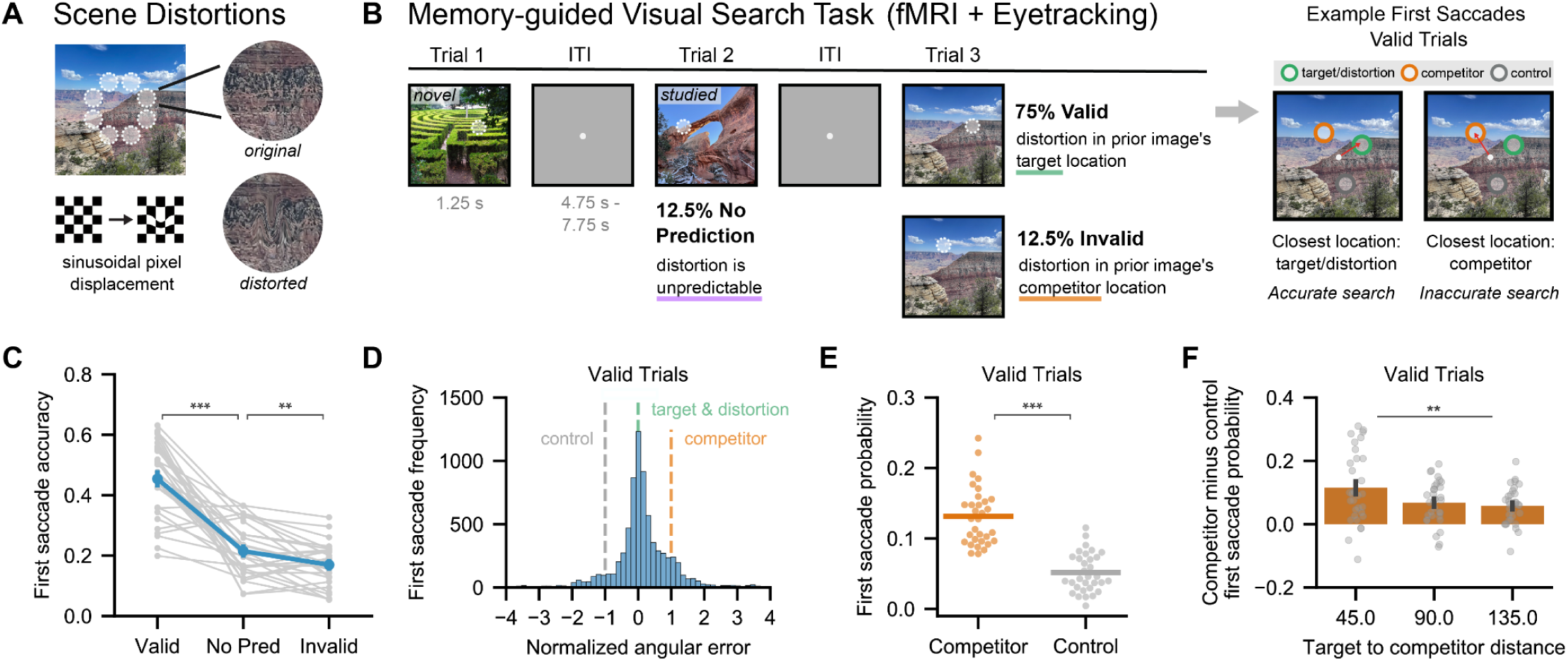
Experimental design and behavioral data from memory-guided visual search task (session 2). **(A)** Scene distortions were created by applying a sinusoidal pixel displacement to a local image patch. **(B)** Memory-guided visual search task. Participants located each distortion (white circle, visualization only) via eye movements. Distortion locations were most likely to be the previous image’s target location (valid trials: 75%). Distortions could also appear in the competitor location (invalid trials: 12.5%), or were unpredictable following novel images (no-prediction trials: 12.5%). Saccade endpoints were evaluated relative to the distortion, target, competitor, and an equidistant control location. Example saccades for a valid trial are shown at right. **(C)** Relative to no-prediction trials, participants’ first saccade was more accurate on valid trials (t(31) = 8.72, p < 0.001, d_z_ = 2.30, valid - no-prediction = 0.24, 95% CI = [0.18, 0.30]) and less accurate on invalid trials (t(31) = −3.44, p = 0.002, d_z_ = 0.60, invalid - no-prediction = −0.05, 95% CI = [−0.07, −0.02]), demonstrating memory-guided search (N = 32 participants, two-tailed paired t-tests). Gray lines indicate individual participants; blue lines indicate across-participant mean ± SEM. **(D)** First saccade angular error distribution on valid trials. Errors are normalized by target-competitor distance (x=0: target/distortion; x=1: competitor; x=−1: equidistant control). Saccades were most frequent near the target, but the distribution is asymmetric toward the competitor. **(E)** First saccade probabilities for competitor and control locations from (D). Saccades to the competitor were more frequent than to the control (t(31) = 7.59, p < 0.001, d_z_ = 2.25, competitor − control = 0.08, 95% CI = [0.06, 0.10]; N = 32 participants, two-tailed paired t-test), suggesting that competition persisted into the search task. Dots indicate individual participants; lines indicate across-participant means. **(F)** Target-competitor distance modulated first saccade likelihood to the competitor relative to control (F(2,62) = 5.93, p = 0.004, η^2^_P_ = 0.16; N = 32 participants; repeated measures ANOVA). Errors were highest when locations were closest. Bars indicate across-participant mean ± SEM. ** p < 0.01; *** p < 0.001.

To test whether participants were using memory to improve search performance, we examined the accuracy of participants’ saccades across the three conditions. Similar to standard attentional cueing effects, we expected participants to perform best in the valid condition, when their predictions were accurate, and worst in the invalid condition, when their predictions were systematically wrong. To verify this, we calculated the angular error between participants’ first saccade end point and the center of the distortion location on each trial and scored this saccade as accurate if it was closer to the distortion than to any of the other 7 possible locations (<22.5 degrees of angular error). We expected the first saccade of the search interval to show the strongest evidence for a memory-based strategy if participants were using the previous scene cue to prepare a saccade. As predicted, we found highly reliable differences in accuracy across the three conditions (F(2,62) = 78.7, p < 0.001, η^2^_P_ = 0.72, Fig. 2C). Participants were more likely to make accurate first saccades on valid trials compared to no-prediction trials (t(31) = 8.72, p < 0.001, d_z_ = 2.30, valid - no-prediction = 0.24, 95% CI = [0.18, 0.30]) and invalid trials (t(31) = 9.74, p < 0.001, d_z_ = 2.91, valid - invalid = 0.28, 95% CI = [0.23, 0.34]). Participants were also less likely to make accurate first saccades on invalid trials than on no-prediction trials (t(31) = −3.44, p = 0.002, d_z_ = 0.60, invalid - no-prediction = −0.05, 95% CI = [−0.07, −0.02]). These effects are consistent with previously reported attentional cueing effects, including memory-based attentional cueing^19^; they verify that participants prepared a saccade to the target location, yielding performance benefits when the distortion appeared there and performance decrements when it unexpectedly appeared elsewhere. We next examined the final saccade of the search interval to test whether these effects persisted into the trial. Relative to the first saccade, the final saccade of the search interval was more accurate in every condition (Supplementary Fig. 1A). However, condition differences were still highly significant for the final saccade (F(2,62) = 65.9, p < 0.001, η^2^_P_ = 0.68 with valid performance highest and invalid performance lowest (all pairwise comparisons significant at p < 0.001 and d_z_ > 0.61; Supplementary Fig. 1A). These results establish that participants were using memory for the scene-locations associations to guide visual search.

To determine whether memory competition was present during visual search, we evaluated whether participants sometimes used the competing memory to guide their attention. To that end, we looked at the distribution of first saccade locations on valid trials. Because the distortion was always located at the target location on these trials, participants had no reason to look at the competitor location first unless they had retrieved the wrong memory. As expected, the first saccade on these trials was frequently near the target (and distortion) location (Fig. 2D and Supplementary Fig. 2). Critically, more first saccades were close to the competitor location than to a control location equally far from the target (t(31) = 7.59, p < 0.001, d_z_ = 2.25, competitor – control = 0.08, 95% CI = [0.06, 0.10]; Fig. 2E and Supplementary Fig. 2). Because it was not behaviorally advantageous to make a first saccade to the competitor location, which was far less likely to contain the distortion than the target location, this pattern of behavior suggests that interference between the similar scenes created competition in attention and memory. Relative to the first saccade, participants had a reduced tendency to look at the competitor location on the final saccade of the search (Supplementary Fig. 1B and 1C). However, final saccades to the competitor location were still more frequent than to the control location, indicating persistent memory competition over the trial (t(31) = 5.05, p < 0.001, d_z_ = 1.22, competitor - control = 0.035; 95% CI = [0.02, 0.05]; Supplementary Fig. 1C).

Given these results, we next investigated whether memory competition was modulated by the distance between target and competitor locations. We predicted that higher similarity (smaller distance) between target and competitor locations would yield more competition based on prior findings that similarity influences memory interference^33^. Consistent with this, we found the largest amount of competition when the target and competitor locations were adjacent along the circle of possible locations (45 degrees apart). Specifically, participants made relatively more first saccades to the competitor location than to a distance-matched control location when the target and competitor locations were 45 degrees apart than when they were 90 degrees or 135 degrees apart (F(2,62) = 5.93, p = 0.004, η^2^_P_ = 0.16; Fig. 2F and Supplementary Fig. 2). There were no reliable differences between target-competitor location distances when examining the last saccade (F(2,62) = 0.265, p = 0.768, η^2^_P_ = 0.008; Supplementary Fig. 1D), though note the low overall number of final saccades to the competitor location. Together, these results confirm that participants performed accurately on the memory-guided search task but nevertheless experienced memory competition.

Because there was a clear effect of memory competition in our task, we next investigated whether participants were systematically biased toward or away from the competitor location. In order to test this in a way that was independent of our previous analyses detecting interference errors, we looked for bias within correct responses (<22.5 degrees of error). Indeed, using a signed measure of first saccade error (positive values toward the competitor), we found that correct saccades were reliably biased toward the competitor (t(31) = 2.43, p = 0.021; d_z_ = 0.43, error = 0.81 degrees toward competitor, 95% CI = [0.13, 1.49]). Although a bias toward the competitor location is consistent with memory competition, this effect was small, and an opposite bias away from competitor features has also been reported^34^. Thus, we do not over-interpret this effect, which is more ambiguous than the error-based analyses presented earlier.

Finally, though our manipulations of interest were within-participants, we next explored individual differences in search performance. We expected that individual differences in search performance would be predicted by individual differences in memory if participants were relying on memory to guide search. To investigate this, we calculated the correlation between search performance on valid trials in session 2 and final memory performance in session 1. Across-participant variance in search performance was strongly predicted by variance in final memory performance in session 1 (proportion of saccades near target location in session 1 vs. session 2: r(30) = 0.56, p < 0.001, 95% CI = [0.26, 0.76]). Furthermore, the degree of memory competition experienced by participants in session 1 predicted competition on valid trials in session 2 (proportion of saccades near competitor location in session 1 vs. session 2: r(30) = 0.40, p = 0.025, 95% CI = [0.05, 0.65]), indicating that saccades to the competitor location in session 2 were most likely the result of memory competition. These results further validate the importance of memory, and the resolution of memory competition, for visual search performance.

### Distinct hippocampal representations of competing scenes predict search behavior

We focused our fMRI analyses of the memory-guided search task on two regions that were of strong theoretical interest: 1) the hippocampus, which is critical for resolving memory competition^16,35,36^, and 2) early visual cortex, whose representations are biased in favor of attentional targets^37,4,5^. For each participant, we defined the hippocampus (HIPP) anatomically in native space. To define early visual cortex (EVC), we combined areas V1, V2, and V3, each of which were defined functionally in native space using data from a separate retinotopic mapping task (see Methods/Regions of Interest). We also defined several regions of interest (ROIs) to serve as controls. These included the dorsal attention network (DAN) and several scene-selective regions: the parahippocampal cortex (PHC), the parahippocampal place area (PPA), and retrosplenial cortex (RSC). For details on how these were defined, see Methods/Regions of Interest.

First, we sought to link the separability of competing scene representations in the hippocampus to memory-guided search behavior. Following prior work^13,16^, we computed an fMRI pattern similarity metric of scene separability. We first estimated the average pattern of BOLD activity evoked by every scene image in the search task (24 paired scenes and 56 novel scenes; see Methods/Pattern Similarity Analyses). For each of the 12 scene pairs, we computed the neural pattern similarity (z-transformed Pearson correlation) of the pairmate images (e.g. arch 1 and arch 2, pairmate pattern similarity; Fig. 3A). For each scene pair, we also computed the average pattern similarity between each of the pairmates and all non-pairmate images (e.g. arch 1 and barn 2, nonpairmate pattern similarity). Pattern similarity with novel images was excluded from the nonpairmate average to control for image familiarity. For each scene pair, we then computed the difference between pairmate and nonpairmate pattern similarity, which we refer to as scene pair pattern similarity. Critically, this difference score indicates the degree to which pairmates are distinct in a given brain region, controlling for the average similarity across images in that region. Higher values indicate that pairmates are relatively more similar to each other than they are to other images, whereas lower values indicate that pairmates are relatively more distinct from each other compared to other images. Similar to prior fMRI studies on hippocampal differentiation^13,16^, we observed lower scene pair pattern similarity values in the hippocampus than in visual cortex (t(31) = −10.2, p < 0.001, d_z_ = 2.33, HIPP - EVC = −0.041, 95% CI = [−0.05, −0.03]; Fig. 3B).

**Figure 3.**
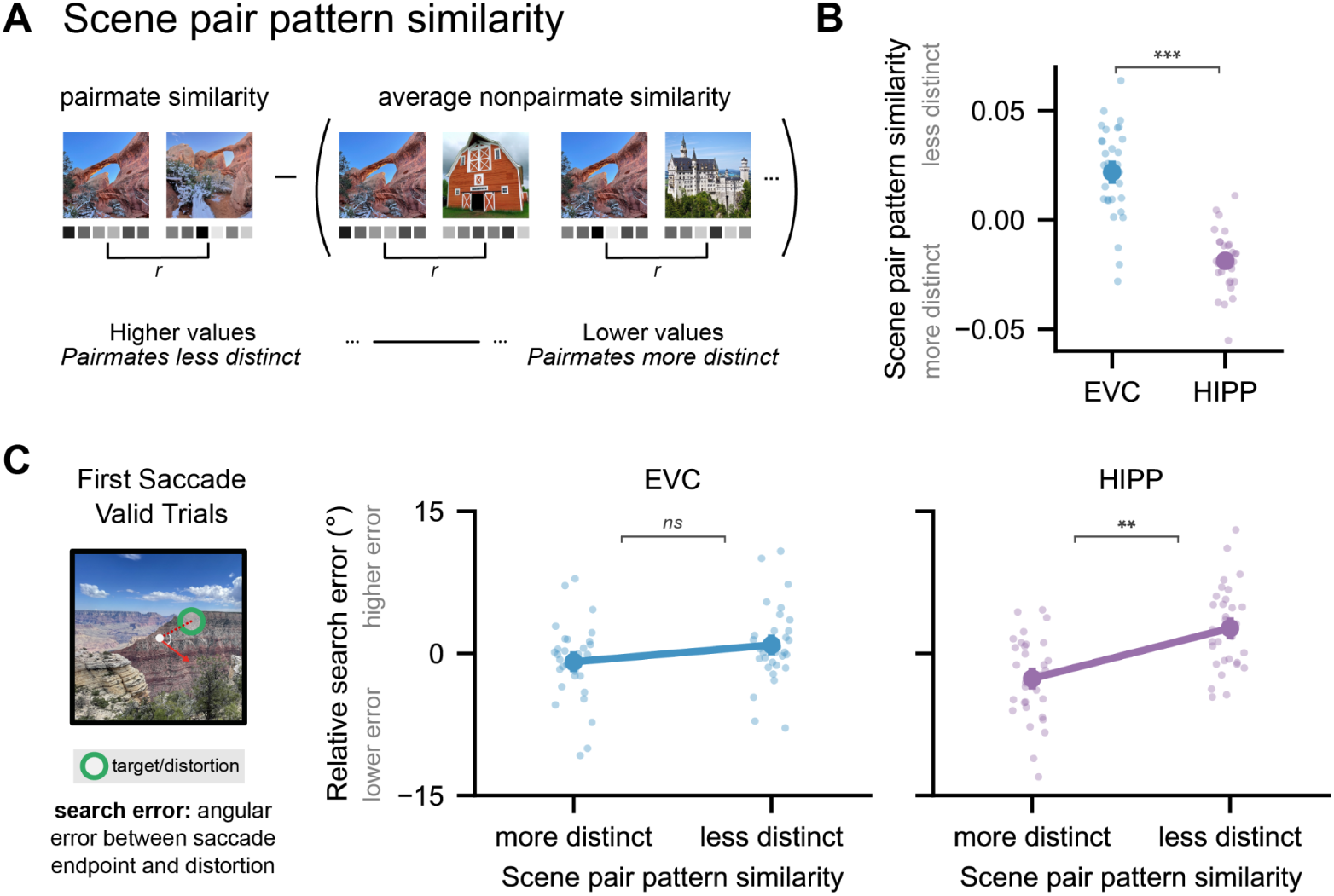
Distinct hippocampal representations of competing scenes predict memory-guided eye movements. **(A)** For every pair of similar scenes, we computed scene pair pattern similarity as the difference between pairmate similarity (e.g. arch 1 and arch 2) and average nonpairmate similarity (e.g. arch 1 and barn 2). Under this measure, lower values indicate that pairmates were relatively more distinct from each other compared to other scenes. **(B)** We compared average scene pair pattern similarity in early visual cortex (EVC; defined as V1, V2, and V3, indicated in blue) and the hippocampus (HIPP, indicated in purple). Consistent with prior work, competing scenes were more distinct in the hippocampus relative to visual cortex (t(31) = −10.2, p < 0.001, d_z_ = 2.33, HIPP - EVC = −0.041, 95% CI = [−0.05, −0.03]; N = 32 participants, two-tailed paired t-test). Small dots represent individual participants. Large dots represent the across-participant mean ± SEM. **(C)** Scene pair pattern similarity in the hippocampus predicted memory-guided eye movements on valid trials. Scene pairs that were more distinct in the hippocampus were followed by first saccades that were closer to the distortion location (also the target location) on the next trial (t(31) = −3.46, p = 0.002, d_z_ = 0.32, more distinct - less distinct = −5.3 degrees, 95% CI = [−8.43, −2.18]; N = 32 participants, two-tailed paired t-test). There was no significant relationship in EVC (t(31) = −1.24, p = 0.223, d_z_ = 0.11, more distinct - less distinct = −1.78 degrees, 95% CI = [−4.69, 1.14]). Error values (angular error between first saccade endpoint and the distortion on valid trials) are centered within participants for visualization but see Supplementary Figure 3A for uncentered data. Small dots represent individual participants. Large points represent the across-participant mean ± SEM. See Supplementary Figure 3 and 4 for related analyses. ns = p > 0.05; ** p < 0.01; *** p < 0.001.

Critically, we predicted that variability in this metric would be related to memory-guided search behavior on trials in which memory was helpful (valid trials); and, further, that this brain-behavior coupling would be selective to the hippocampus. Because our task required discrimination between competing scenes to make an accurate memory-guided saccade on the next trial, we hypothesized that more separable memory representations in the hippocampus would be associated with saccades that were closer to the distortion location in the next search period. We reasoned that the first saccade of the search period would most strongly reflect memory processing because it could be planned prior to stimulus presentation when no bottom-up input was present. Thus, for every participant and scene pair, we calculated the average error between the first saccade endpoint and the distortion location on valid trials following the presentation of either scene pairmate of interest. Indeed, scene pairs that were more distinct in the hippocampus (i.e. with lower scene pair pattern similarity) were followed by more accurate (lower error) saccades than scene pairs that were less distinct (t(31) = −3.46, p = 0.002, d_z_ = 0.32, more distinct - less distinct = −5.3 degrees, 95% CI = [−8.43, −2.18]; Fig. 3C, right and Supplementary Fig. 3A). To test whether this relationship was robust to different behavioral metrics and analysis choices, we performed this analysis several other ways. First, we tested whether this relationship was present when we used the binarized saccade accuracy measure previously reported in Figure 2 (<22.5 degrees defined to be accurate) rather than a continuous measure of error. Scene pairs that were more distinct in the hippocampus were followed by more accurate saccades under this measure (t(31) = 2.04, p = 0.049, d_z_ = 0.21, more distinct - less distinct = 0.026, 95% CI = [0.0, 0.05]; Supplementary Fig. 3B). This effect became stronger when using a more stringent accuracy threshold of 10 degrees (t(31) = 2.67, p = 0.012, d_z_ = 0.31, more distinct - less distinct = 0.021, 95% CI = [0.0, 0.04]). We also confirmed that the results from the continuous error analysis were consistent when analyzing the last saccade instead of the first saccade (t(31) = −3.01, p = 0.005, d_z_ = 0.23, more distinct - less distinct = −3.4 degrees, 95% CI = [−5.73, −1.10]; Supplementary Fig. 3C), and when allowing scene pair similarity scores to vary continuously rather than binning them into high and low groups (t(315) = 2.38, p = 0.018, β = 1.98, 95% CI = [0.34, 3.62]; Supplementary Fig. 3D). Finally, we confirmed that scene pair pattern similarity in the hippocampus was not significantly related to the number of saccades participants made when viewing the scenes (t(31.0) = −0.06, p = 0.949, β = −0.001, 95% CI = [−0.04, 0.04]).

We next performed an exploratory analysis to assess whether the relationship between hippocampal representations and behavior varied across its anterior-posterior axis, given prior work showing important anatomical and functional differences between anterior and posterior hippocampus^38^. We found a significant interaction between ROI (anterior/posterior hippocampus) and scene pair pattern similarity on search error (F(1,31) = 10.76, p = 0.003, η^2^_P = 0.26_) such that more distinct representations for scene pairmates were linked to lower search error in anterior (t(31) = −4.33, p < 0.001, d_z_ = 0.43, more distinct - less distinct = −7.1 degrees, 95% CI = [−10.44, −3.75]) but not posterior (t(31) = −0.11, p = 0.915, d_z_ = 0.01, more distinct - less distinct = −0.19 degrees, 95% CI = [−3.84, 3.45]; Supplementary Fig. 4A,B) hippocampus. Thus, while our study was not designed to test this question, we observed that anterior hippocampal representations were selectively related to more accurate and interference-free memory-guided eye movements.

Finally, we sought to test whether this relationship was selective to the hippocampus. We first performed the same analysis in several visual cortical ROIs with sensitivity to visual stimuli and/or scenes (EVC, PPA, RSC, and PHC). Considering all five ROIs together – i.e., including hippocampus – there was a significant interaction between ROI and scene pair pattern similarity bin (F(4,124) = 2.57, p = 0.041, η^2^_P_ = 0.077), indicating that the relationship between representations and behavior varied across these brain areas. In contrast to the hippocampus, there was no statistically significant relationship between scene pair pattern similarity in EVC and saccade error (t(31) = −1.24, p = 0.223, d_z_ = 0.11, more distinct - less distinct = −1.78 degrees, 95% CI = [−4.69, 1.14]; Fig. 3C, left). Similarly, there was no relationship in PPA (t(31) = −0.14, p = 0.890, d_z_ = 0.017, more distinct - less distinct = −0.28 degrees, 95% CI = [−4.36, 3.80]), RSC (t(31) = 0.90, p = 0.373, d_z_ = 0.098, more distinct - less distinct = 1.63 degrees, 95% CI = [−2.05, 5.31]), or PHC (t(31) = −1.32, p = 0.198, d_z_ = 0.15, more distinct - less distinct = −2.55 degrees, 95% CI = [−6.51, 1.41]; Supplementary Fig. 4C,D). To further investigate selectivity to the hippocampus, we performed an exploratory whole-brain searchlight analysis to look for regions in which search error was predicted by scene pair pattern similarity (more vs. less distinct pairmates, following the procedure in Fig. 3). This analysis revealed clusters in the anterior hippocampus and lateral occipital lobe, in which more distinct representations for pairmates was associated with reduced search error (Supplementary Fig. 4E). No other regions showed a relationship between scene pair pattern similarity and search error. Taken together, these results show an important link between hippocampal representations and rapid attentional behavior guided by memory, and raise the question of how hippocampal representations are coordinated with visual cortical representations to support attention.

### Preparatory target and competitor location coding in visual cortex

We next sought to understand how retrieved memories were used to benefit visual search. Our search task was designed so that participants could use memory to predict the location of the upcoming distortion during the blank inter-trial interval (ITI) on the vast majority of trials (valid trials). Because early visual cortex contains maps of the visual field, and because these maps can be activated in a top-down manner^39–42^, we hypothesized that participants’ location predictions during the ITI would be represented by spatially-organized activity within these maps. We refer to such spatial representations during the ITI as preparatory coding because they occur prior to the onset of the stimulus and are in direct support of the upcoming task. Given how critical memory-based prediction was to task performance (see Fig. 2C), we hypothesized that preparatory target location coding in visual cortex would be associated with search success on the next trial, whereas preparatory competitor location coding would be associated with interference errors.

To test these hypotheses, we took advantage of models that were developed to characterize the spatial organization of visually-evoked fMRI signals. Using data from a separate retinotopic mapping task, we estimated the best-fitting population receptive field (pRF) for each voxel in a participant’s brain (Fig. 4A and see Methods/pRF Model Fitting). This pRF represents the location and extent in the visual field of that voxel’s maximal response to visual stimulation. We used the pRF parameters to isolate voxels in early visual cortex (EVC; defined as V1, V2, and V3) with responsiveness to the 8 possible target locations. We then examined the BOLD response in these EVC voxels during the blank ITI of the search task, after regressing out task-evoked activity to remove confounding effects of the visual response from the preceding trial (Fig. 4A and see Methods/Visual Cortex Preparatory Responses).

**Figure 4.**
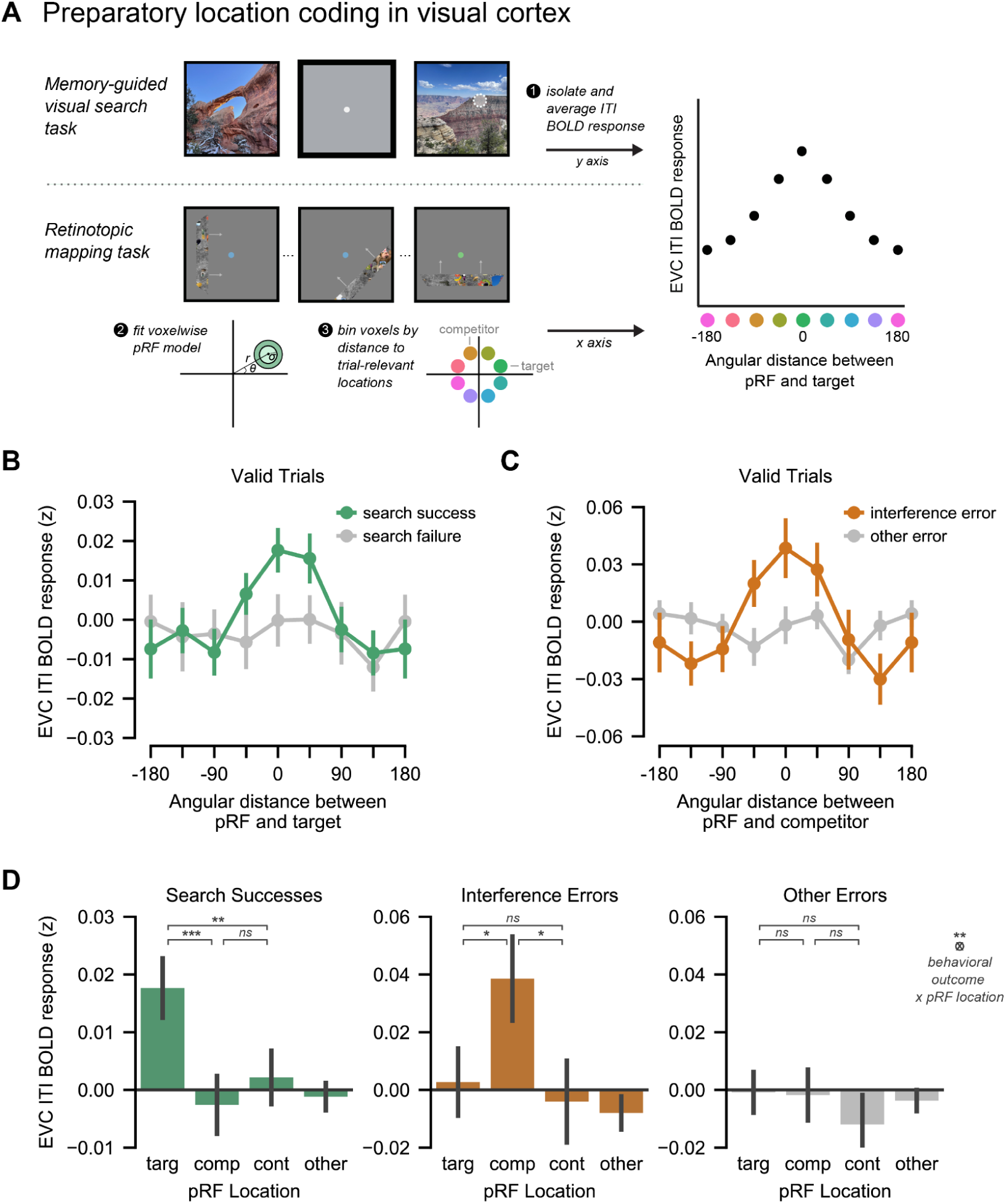
Preparatory coding in early visual cortex predicts search success and failures due to interference. **(A)** During the blank inter-trial interval (ITI) of the search task, participants maintained fixation while engaging in memory-based prediction, allowing examination of spatially organized preparatory activity in early visual cortex (EVC). EVC population receptive field (pRF) parameters were estimated using a retinotopic mapping task: *r* = eccentricity, *θ* = polar angle, *σ* = size. BOLD responses during the ITI were plotted for voxels whose pRFs varied in distance from the upcoming target or competitor locations. **(B)** Preparatory target location coding on valid trials, compared for upcoming search successes and failures, defined by final saccade location (N = 32 participants). Preparatory activity at the target location was higher during ITIs preceding search successes than search failures. Points indicate across-participant mean ± SEM. Data at ±180 degrees are identical and redundantly plotted for visualization. **(C)** Preparatory competitor location coding compared for upcoming interference errors versus other search failures (N = 32 participants). Preparatory activity around the competitor location was higher during ITIs preceding interference errors than other errors. Plotting conventions as in (B). **(D)** We directly compared EVC BOLD activity during the ITI at pRFs corresponding to the target location (targ), competitor location (comp), control location (cont), and all other locations (other) for three behavioral outcomes: search successes, interference errors, and other errors. A linear mixed-effects model revealed an interaction between pRF location and behavioral outcome (F(6,75101) = 3.12, p = 0.005; N = 75144 trials across 32 participants; Type III F-test). This interaction was driven by preferential target activation for search successes, preferential competitor activation for interference errors, and no significant preferential activation for other errors. Only pairwise statistical comparisons between target, competitor, and control locations are shown. Bars represent the across-participant mean ± SEM. Individual participant data can be found in Supplementary Figure 5A. ns = p > 0.05; *p < 0.05; **p < 0.01; *** p < 0.001.

To investigate whether there was a link between preparatory coding of the target location and search success, we first visualized ITI responses as a function of a voxel’s distance between its pRF and the target memory location on the upcoming trial. We did this separately for upcoming trials that were search successes vs. search failures, defined according to the participant’s final saccade. We focused on valid trials (75% of all trials), because these were the trials in which memory yielded accurate predictions. Thus, search successes were trials in which the search concluded at the distortion (also the target memory location) and search failures were all other trials. Consistent with our hypothesis, visualizations of visual cortex representations differed for search successes and failures (Fig. 4B). For upcoming search successes, we observed the strongest ITI response in voxels with pRFs closest to the target location, and this response became weaker as pRFs got further away. For upcoming search failures, this visualization did not reveal differential coding of the target (vs. other) locations during the ITI.

To visualize the relationship between preparatory coding of the competitor location and interference errors, we performed the same procedure as in the prior analysis, but sorted voxels based on their distance to the competitor location. We focused specifically on search failure trials from the previous analysis, averaging separately for upcoming interference errors and other error trials. Because we focused this set of analyses on valid trials, interference errors were searches that ended at the competitor location, despite the fact that the distortion was in the target location. Consistent with our hypothesis, this visualization suggested different representations for upcoming interference errors vs. other errors (Fig. 4C). For upcoming interference errors, we observed the strongest ITI response in voxels with pRFs closest to the competitor location, and this response became weaker as pRFs got further away. For other errors, this visualization did not reveal differential coding of the competitor (vs. other) locations during the ITI.

To quantify these effects simultaneously, each pRF bin was assigned one of the following labels separately for every trial and participant: target, competitor, control, or other. ITI responses in these voxel groups were then compared across behavioral outcomes (search successes, interference errors, and other errors) using a linear mixed effects model with participant as a random intercept (see Methods/Statistics). The interaction between pRF location and behavioral outcome was significant (Type III F-test: F(6,75101) = 3.12, p = 0.005), indicating that preparatory activity in visual cortex voxels coding for target, competitor, control, and other locations varied depending on the behavioral outcome of the upcoming trial.

To clarify this interaction, estimated marginal means of ITI responses were examined across pRF locations for each behavioral outcome. We focus on contrasts between the target location and the competitor and control locations, which were equidistant from the target. For search successes, ITI activity in target pRFs was greater than in competitor (t(75101) = 3.96, p < 0.001, β = 0.013, 95% CI = [0.007, 0.020]) and control (t(75101) = 2.99, p = 0.003, β = 0.011, 95% CI = [0.004, 0.018]) pRFs, indicating preferential activation of the target location (Fig. 4D, left). There was no statistically significant difference between competitor and control pRFs (t(75101) = −0.68, p = 0.498, β = −0.002, 95% CI = [−0.009, 0.005]). For interference errors, ITI activity in competitor pRFs was greater than in target (t(75101) = 2.01, p = 0.044, β = 0.020, 95% CI = [0.0005, 0.040]) and control (t(75101) = 2.52, p = 0.012, β = 0.027, 95% CI = [0.006, 0.048]) pRFs, indicating preferential activation of the competitor location (Fig. 4D, middle). There was no statistically significant difference between target and control pRFs (t(75101) = 0.63, p = 0.528, β = 0.007, 95% CI = [−0.014, 0.028]). For other errors, no statistically significant differences were observed across pRF locations (all |t| ≤ 1.33, all p > 0.18; Fig. 4D, right). These results were similar when using the first rather than the last saccade to define behavioral outcomes (Type III F-test: F(6,75101) = 3.03, p = 0.006; Supplementary Fig. 5B).

Overall, these results demonstrate that memory-based predictions lead to spatially precise preparatory representations in early visual cortex. This precise preparatory coding in turn predicts the accuracy of upcoming eye movements during visual search. However, these results also demonstrate that preparatory representations in early visual cortex can also reflect inaccurate predictions that occur as a result of memory competition. In this case, these representations are associated with interference errors in attentional allocation that lead to task failure. Taken together, our findings in visual cortex show that preparatory coding predicts the successful use of memory to guide attention as well as failure caused by competition.

### Distinct hippocampal representations are associated with reduced competitor activation in visual cortex

Next, we asked whether the separability of scene representations in the hippocampus was related to the relative activation of target and competitor locations in visual cortex. We hypothesized that distinct scene representations in the hippocampus would be associated with reduced activation of competitor locations in visual cortex, allowing preparatory representations in this region to be more precise. To test this hypothesis, we used our pRF-based approach to re-examine preparatory target and competitor location coding in visual cortex during the ITI of valid trials, separately for scenes that were more vs. less distinct in the hippocampus (following the procedure in Fig. 3A). We focused on scene pairs for which the target and competitor locations were 45 degrees apart (adjacent locations in our study) because they posed the most memory competition – specifically, a greater tendency to make saccades to the competitor location vs. the distance-matched control location (Fig. 2F). We first visualized ITI activity in early visual cortex as a function of pRF distance from the target location, separately for 45 degree pairs that were more vs. less distinct in the hippocampus. We did this first at the individual participant level, because the clockwise vs. counterclockwise positioning of the competitor and control locations with respect to the target location differed across scene pairs and across participants for a given scene pair (Fig. 5A). Then, as a visualization, we aggregated this data across participants by averaging within each pRF distance bin (Fig. 5B). Note that this averages together data from competitor and control locations, but provides a visualization similar to those in Figure 4B.

**Figure 5.**
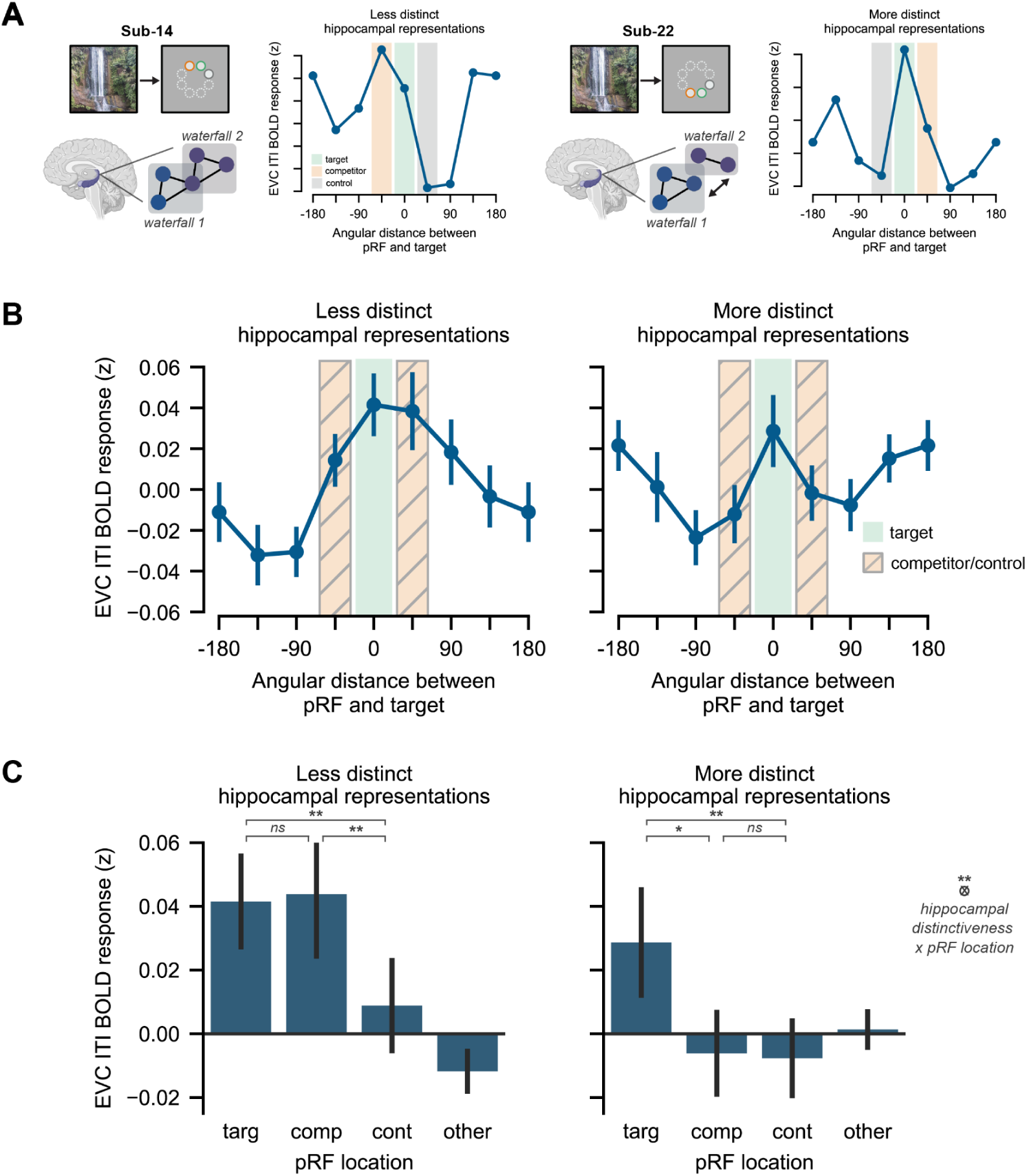
Distinct hippocampal representations are associated with reduced competitor activation in visual cortex. **(A)** Analysis framework for examining the relationship between hippocampal scene representations and preparatory visual cortical coding in scene pairs with adjacent target and competitor locations. Two single-participant early visual cortex (EVC) BOLD responses from the blank intertrial interval (ITI) are plotted, as a function of hippocampal memory distinctiveness for the preceding scenes (low or high). The x axis groups voxels according to population receptive field (pRF) distance from the target, with target (x=0, green shading), competitor (x = +/-45, orange shading), and control (x = +/-45, gray shading) locations indicated. Brain images created in BioRender. Favila, S. (2026) https://BioRender.com/jujv83a. **(B)** Group EVC data averaged across participants (N = 32) within each pRF distance bin. Activity in the ±45 bins (orange/gray shading) includes both competitor and control locations due to randomized spatial positioning across participants and scene pairs. Points indicate across-participant mean ± bootstrapped SEM. Data at ±180 are identical and redundantly plotted. **(C)** Visual cortex ITI responses were quantified by grouping trials into pRF location bins (target, competitor, control, other) and splitting scene pairs by hippocampal memory distinctiveness (more vs. less distinct) within participant. A linear mixed-effects model revealed a significant interaction between pRF location and hippocampal distinctiveness (F(3,19617) = 4.78, p = 0.002; N = 19656 trials across 32 participants; Type III F-test). This interaction was driven by no significant difference between target and competitor activation for less distinct pairs (t(19617) = 0.07, p = 0.943, β = 0.0006, 95% CI = [−0.015, 0.016]), but higher target than competitor activation for more distinct pairs (t(19617) = 2.50, p = 0.013, β = 0.018, 95% CI = [0.004, 0.032]; N = 19656 trials across 32 participants; two-tailed post-hoc contrasts). Only pairwise statistical comparisons between target, competitor, and control locations are shown. Bars represent the across-participant mean ± SEM. Individual participant data can be found in Supplementary Figure 6A. * p < 0.05; ** p < 0.01.

To quantify these effects, we compared ITI responses in target, competitor, control, and other voxels for more vs. less distinct scene pairs (Fig. 5C). We used a linear mixed-effects model to assess the interaction between hippocampal scene distinctiveness (more vs. less) and relative BOLD activity in different pRF locations, with participant included as a random effect (see Methods/Statistics). The interaction between hippocampal distinctiveness and pRF location was significant (Type III F-test: F(3,19617) = 4.78, p = 0.002), indicating that preparatory activity in visual cortex voxels coding for different locations varied depending on hippocampal distinctiveness.

To clarify this interaction, estimated marginal means of ITI responses were examined across pRF locations for each level of hippocampal distinctiveness. We focused on contrasts between the target location and the competitor and control locations, which were equidistant from the target. For scene pairs that were less distinct in the hippocampus, visual cortex ITI activity in target pRFs was not statistically different from competitor pRFs (t(19617) = 0.07, p = 0.943, β = 0.0006, 95% CI = [−0.015, 0.016]). In addition, ITI activity in both target (t(19617) = 2.92, p = 0.004, β = 0.023, 95% CI = [0.007, 0.038]) and competitor (t(19617) = 2.85, p = 0.004, β = 0.022, 95% CI = [0.007, 0.037]) pRFs was greater than in control pRFs (Fig. 5C, left). Thus, when scene pairmates had less distinct representations in the hippocampus, both target and competitor locations were activated in visual cortex but not differentially so. For scene pairs that were more distinct in the hippocampus, ITI activity in target pRFs was higher than in competitor pRFs (t(19617) = 2.50, p = 0.013, β = 0.018, 95% CI = [0.004, 0.032]) and control pRFs (t(19617) = 2.73, p = 0.006, β = 0.020, 95% CI = [0.006, 0.034]). ITI activity in competitor pRFs was not statistically different from control pRFs (t(19617) = 0.24, p = 0.814, β = 0.0017, 95% CI = [−0.012, 0.016]; Fig. 5C, right). Thus, more separable hippocampal scene representations were linked to relative suppression of competitor locations in visual cortex, allowing for more precise visual cortical representations prior to search onset. Complementing our finding that memory competition was elevated for scene pairmates whose target and competitor locations were adjacent, i.e., 45 degrees apart (Fig. 2F), the interaction between hippocampal scene distinctiveness and visual cortex pRF responses was selective to the 45 degree pairs. The same interaction was not statistically significant for 90°, 135°, or 180° pairs (all Fs < 0.91, all ps > 0.40), and the three way interaction between target–competitor distance, pRF location, and hippocampal distinctiveness was significant (Type III F-test: F(6,71597) = 2.59, p = 0.016). The selectivity of this effect is consistent with work proposing that the amount of competition between memories is a driver of representational change in the brain10,43,44.

We next performed an exploratory analysis to assess whether the relationship between hippocampal distinctiveness and visual cortex activation varied across the hippocampal anterior-posterior axis. We considered only the scenes whose target and competitor locations were 45 degrees apart because effects in the whole hippocampus were selective to this condition. There was no interaction between hippocampal long-axis position, scene pair distinctiveness, and visual cortex pRF location (Type III F-test: F(3,39236) = 1.12, p = 0.340). Instead, the results in both anterior (F(3,19588.8) = 2.39, p = 0.066) and posterior (F(3,19591.4) = 2.29, p = 0.077) hippocampus were qualitatively similar to those in the whole hippocampus (Supplementary Fig. 6B).

We then sought to test whether relative target and competitor activation in visual cortex was selectively related to scene distinctiveness in the hippocampus. To test this, we repeated our previous analysis of the 45 degree pairs but divided our visual cortex data using pattern distinctiveness in other brain regions besides the hippocampus. We used the same control regions of interest as we did for Figure 3 (see results in Supplementary Figure 4), and tested whether pattern distinctiveness in these regions (PPA, RSC, and PHC) predicted location coding in early visual cortex (Supplementary Fig. 6C). There was no statistically significant interaction between scene pair distinctiveness in PPA (F(3,19617) = 0.44, p = 0.725) or PHC (F(3,19587.5) = 1.40, p = 0.239) and relative BOLD activity in visual cortex voxels coding for different locations. There was an interaction between scene pair distinctiveness in RSC and relative BOLD activity in visual cortex (F(3,19587.7) = 3.17, p = 0.023). However, the interaction present in RSC differed from the interaction present in the hippocampus and did not easily afford a similar interpretation. First, unlike hippocampus, target and competitor activation did not significantly differ when considering either low (t(19588) = 1.64, p = 0.101, β = 0.012, 95% CI = [−0.002, 0.027]) or high (t(19588) = 1.03, p = 0.304, β = 0.0076, 95% CI = [−0.007, 0.022]) pattern distinctiveness in RSC. Second, the relationship between competitor and control activation was modulated by RSC distinctiveness in the opposite way it was modulated by hippocampal distinctiveness, such that more distinct RSC representations were related to relatively higher competitor than control activation (t(19588) = 3.42, p < 0.001, β = 0.025, 95% CI = [0.011, 0.040]). Finally, we considered whether univariate activity in the dorsal attention network (DAN), a network of regions in frontoparietal cortex implicated in attentional control over the visual system, could be related to relative target and competitor activation in visual cortex. Instead of dividing our data by pattern distinctiveness, we divided scene pairs according to their evoked univariate BOLD activity in the dorsal attention network (DAN). We found no statistically significant relationship between univariate activity in the DAN and relative BOLD activity in visual cortex voxels coding for different locations (F(3,19592.3) = 1.155, p = 0.325; Supplementary Fig. 6C). Taken together, these results demonstrate a selective link between the distinctness of hippocampal representations and preparatory signals in the visual system that support memory-guided search behavior.

## Discussion

Memory and perception both require selection of relevant representations in the face of competitors. Here, using an original task, we demonstrate that mechanisms that overcome competition in memory and perception are linked via the hippocampus. First, we show that more distinct scene representations in the hippocampus are associated with more accurate eye movements during memory-guided visual search. Second, we show that early visual cortex exhibits preparatory coding of target and competing search locations, with preferential representations of upcoming target locations predicting search successes and preferential representations of upcoming competitor locations predicting search failures due to interference.

Finally, we link these findings by showing that, when competition is high, distinct representations in the hippocampus are associated with changes to the precision of preparatory representations in visual cortex. When scene representations in the hippocampus are more distinct, relative competitor activation in visual cortex is lower, yielding more precise preparatory representations.

Our findings build on a rich literature of prior work examining the role of memory in guiding visual attention. Behavioral studies have repeatedly shown that attentional performance, such as the speed of visual search, improves with previous exposure to the search targets and/or surrounding context^45–48^. More recent work has illuminated the circumstances under which memories may cause interference or competition in attentional allocation and eye movements^49–51^. Complementary lesion studies^17,52,22^ and fMRI experiments^19,18,26,53,27^ have also begun to uncover the hippocampal and cortical mechanisms by which memories can guide attention. We were particularly inspired by prior fMRI studies showing that: 1) hippocampal activity is higher when attention is guided by memory than when it is guided by external cues ^19,27^ and 2) contralateral preparatory activity in visual cortex precedes memory-guided attention^26^.

Here, we sought to extend prior work implicating the hippocampus in memory-guided attention by 1) testing how hippocampal memory distinctiveness shapes the precision of attentional selection; and 2) linking these hippocampal representations to spatially organized preparatory coding in visual cortex – an index of spatial memory retrieval in anticipation of upcoming attentional goals. To do this, we capitalized on several fMRI approaches that allow characterization of representations in the brain: pattern similarity techniques and visual encoding models. These approaches allowed us to test specific hypotheses about how the distinctiveness of hippocampal representations may be linked to the suppression of competing locations in visual cortex.

Our work critically extends prior work on memory-guided attention to more realistic task contexts in which multiple memories compete for attention, revealing ties between how competition is resolved in memory and perception. To make this advance, we drew on recent fMRI studies examining how the hippocampus prevents interference between competing memories^13,14,16^. These studies associated highly similar scenes with distinct outcomes and tracked how hippocampal representations of the scenes changed as memory interference was resolved. This work has found that while similar memories begin with similar hippocampal representations, these representations become differentiated with repeated competition, consistent with computational models that have proposed activity-dependent representational change in the hippocampus^9,10,43^. Complementary work with MEG has shown that the hippocampus may minimize competition between overlapping memories by representing them at different phases of its ongoing theta rhythm^54^. We built on this set of findings by asking whether distinct hippocampal representations support online perceptual behavior and are linked to representations in the visual system. Instead of focusing on the circumstances that lead to hippocampal differentiation or the absolute magnitude of differentiation, as prior work has done^13,14,16^, we focused on variability in hippocampal representations at the end of this process.

Specifically, we reasoned that this variability would be linked to variation in visual cortex representations and in eye movements during visual search. By showing these links, our study demonstrates that memory distinctiveness in the hippocampus is not just important for direct assessments of memory, but also for natural perceptual behaviors in which memory must be combined with ongoing sensory input. This finding adds to the growing literature suggesting that the separability of hippocampal representations is important for diverse cognitive functions, including navigation^55^, control^56^, and reinforcement learning^57^.

Note that because our purpose was not to re-establish prior findings of hippocampal differentiation, we did not measure neural representations of our stimuli prior to learning, nor did we include a baseline condition consisting of similar scene pairs that had no competing associations. Thus, unlike prior studies, our data cannot provide direct evidence for whether similar memories that compete are differentiated in the hippocampus above and beyond similar memories that don’t compete. However, none of our effects of interest are concerned with the overall magnitude of pattern similarity for pairmates vs. nonpairmates (whether these difference scores are above or below zero or by how much). Instead, our focus was on how variability in this measure is related to behavior and representations in visual cortex. Nevertheless, because of prior evidence linking memory competition to changes in hippocampal representations^13,16^ and the similarities between our study and these prior studies, we still interpret our findings in terms of the competition-dependent memory differentiation framework. An alternative explanation is that some scene pairs are represented more distinctly than others even on initial exposure, due to perceptual and/or semantic attributes of the scenes or due to variation in automatic hippocampal pattern separation^58^. This source of variation in hippocampal scene separability, however, cannot account for our findings without also positing additional links between scene representations and learning in our task. For example, scene pairs that are more effectively separated by the hippocampus on initial exposure could support more effective scene-location learning and thus more accurate memory-guided eye movements. Given the prior research showing changes in hippocampal representations due to memory competition^13,16^ – research using similar learning tasks and stimuli as used here – we think it is likely that the competition framework will be supported over one based on automatic pattern separation. Nevertheless, to adjudicate between these accounts, future studies will need to manipulate stimulus properties of interest while also tracking hippocampal representations during initial exposure and over learning.

Stimulus properties may also play a role in explaining our exploratory results comparing the anterior and posterior hippocampus. Though we did not find reliable differences between the anterior and posterior hippocampus in their relationship to preparatory coding in visual cortex, we did find differences in their relationship with search behavior. Post-hoc analyses revealed that the relationship between hippocampal scene distinctiveness and search behavior was selective to the anterior hippocampus. At first pass, these results are at odds with prior work that has proposed that the posterior hippocampus contains detailed representations while the anterior hippocampus contains more gist-like representation^38^. However, experimental evidence in support of this idea has been mixed. Some studies have found evidence for more distinct representations in anterior hippocampus^59^, particularly after a delay^60^. Some studies also report differences in stimulus selectivity along the hippocampal long axis, which may interact with representational granularity accounts. For example, some prior work has argued that the anterior hippocampus plays a particularly important role in scene processing and recall^61^. Thus, further examination of anterior-posterior differences in hippocampal coding, and how they interact with other experimental factors, such as memory delay or stimulus properties, is required.

Our results are consistent with a large body of work demonstrating memory reactivation and preparatory coding in early visual cortex^62,26,63–65,42,66^. Though memory reactivation is a widely documented phenomenon, we extend this body of work in several ways. First, we provide evidence for reactivation of multiple competing memories in early visual cortex. Prior work looking at competitive memory reactivation has primarily relied on competing memories from different categories (e.g., faces and places) and has used diminished category decoding accuracy during recall as an indirect indicator that multiple memories are being reactivated^67,68^. Here, we characterized competitive reactivation in the spatial domain, using model-based methods that distinguish between co-activation of multiple memories and failed retrieval. Second, we show that relative activation between two competing memories predicts the location of upcoming eye movements. This finding is consistent with previous work that reported a link between coarse (hemifield-level) memory reactivation and visual search behavior^26^. We extend this link with behavior to a case in which multiple memories compete and also use methods that allow us to characterize neural reactivation at a higher level of spatial precision.

Finally, we show that hippocampal pattern distinctiveness is selectively related to the precision of visual cortex preparatory coding when memory competition is especially high. Specifically, scene pairmates with a 45 degree target-competitor distance were associated with the highest number of interference errors, suggesting elevated memory competition and raising the possibility that hippocampal distinctiveness may be particularly important for the 45-degree condition. Indeed, when target and competitor locations were 45 degrees apart, more distinct hippocampal representations for the pairmates was associated with preferential activation of the target location in visual cortex. Conversely, less distinct hippocampal representations were associated with activation of both target and competitor locations in visual cortex. This relationship between hippocampal pattern distinctiveness and visual cortex preparatory activity was not present when target and competitor locations were further apart, suggesting that hippocampal distinctiveness may play a particularly important role in modulating visual cortex activity when memory competition remains high. Studies that track the relationship between hippocampal patterns and visual cortex preparatory activity as memory competition is resolved will be needed to confirm this account. The selectivity of the hippocampal-visual cortex relationship to 45 degree target-competitor distances is also consistent with the idea that the hippocampus is particularly important for binding precise, high-resolution information in memory^69,70^. Under this account, the hippocampus may be particularly important in modulating visual cortex when resolving competition between memories requires fine spatial discrimination.

While our primary analyses focused on target, competitor, and control locations, we also observed an elevated preparatory signal 180 degrees from the target location for scene pairs that were more distinctive in the hippocampus (Figure 5B, right). Exploratory analyses revealed that this signal was primarily driven by early visual cortex voxels with low eccentricity pRFs. Participants rarely made eye movements toward the 180 degree location (SFig. 2A), suggesting that the signal arising from these voxels during the ITI is unlikely to correspond to overt behavior in the upcoming search task. Importantly, when we removed voxels with low eccentricity pRFs from our analysis, it reduced the 180 degree signal, but our primary interaction between hippocampal scene distinctiveness and pRF (target, competitor, control, and other) locations remained intact (Type III F-test: F_3, 19617_ = 3.73, p = 0.011). This 180 degree signal may partially be explained by errors in pRF estimation near the fovea, where small positional differences in estimated pRF centers can lead to large differences in the polar angle estimate. However, this would not explain why the signal is only present for scenes pairs that were more distinctive in the hippocampus. One speculative possibility is that some of this signal reflects a form of representational remapping, similar to what has been observed in working memory studies. In these studies, unprioritized items are remapped to a different portion of the representational space in order to decrease competition with prioritized items^71^. In this framework, the enhanced signal at 180 degrees may minimize interference by maximizing the distance between the target and competitor in visual cortex. The specificity of this finding to scene pairs that were more distinct in the hippocampus points to the possibility that hippocampal mechanisms might contribute to remapped representations in visual cortex, an intriguing idea that requires further investigation.

Our results raise the possibility that hippocampal computations may drive precise memory-guided eye movements and influence visual cortical representations. However, it is important to note that fMRI cannot establish causal links between hippocampus, visual cortex, and the oculomotor system. Our hypothesized mechanism is supported by strong behavioral evidence that memory is contributing to behavior in our task, as well as prior work implicating the hippocampus in memory-guided attention^47,31,27,29^ and prediction^72–74,27,75^. Our proposed mechanism is also supported by prior causal studies showing that hippocampal damage disrupts anticipatory coding in visual cortex^76^ and that hippocampal stimulation can drive activity in oculomotor areas^28^. Future studies with stimulation techniques or lesion methods, however, will be critical for direct tests of our hypothesis about causal links between hippocampus, visual cortex, and subsequent eye movements during memory-guided attention. That said, even if such tests determined that the hippocampus can exert a causal influence on visual cortex representations and eye movements, it is likely that interactions between hippocampus and visual cortex are bidirectional. For example, attentional modulation of visual cortex representations could contribute to the distinctiveness of hippocampal representations by adjusting input to the hippocampus^77^.

It is possible that other brain areas are also supplying top-down drive to visual cortex and/or the hippocampus. Candidate areas for such modulation include regions of prefrontal and parietal cortex that are implicated in the control of visual attention^78^, memory-guided attention^79,80,29^, and resolving memory competition^81,82^. One intriguing possibility is that control-related cortical regions play an important role in enabling top-down spatial attention in the face of memory competition early in learning when hippocampal memories are not well formed or too overlapping to prevent interference. However, once hippocampal memory representations are sufficiently differentiated, the need for control resources may be reduced^83^. While we did not observe any link between univariate activity in the dorsal attention network (which includes lateral frontoparietal areas) and visual cortex preparatory coding in this study, it’s important to note that our measurements were made after considerable training occurred in session 1. Future work could examine different neocortical systems as memories are acquired to better understand the contributions of multiple brain areas to competition resolution and memory-guided attention.

To conclude, we show that hippocampal mechanisms for distinguishing competing memories are associated with more accurate attentional selection and with more precise preparatory activity in the visual system. Our results demonstrate linked mechanisms that overcome competition in memory and perception and argue for a broad role for memory systems in shaping adaptive perceptual behavior.

## Methods

### Participants

Thirty-six human participants were recruited from the Columbia University community and the surrounding area. No statistical method was used to predetermine sample size. Participants provided written informed consent prior to the experiment and were compensated $20/hr for their time. All experimental procedures were approved by the Columbia University Institutional Review Board. We excluded all data acquired from four participants from our analyses. Three participants were excluded because they withdrew from the experiment during the first experimental session due to difficulty with the task. MRI data was not acquired for these participants. One participant was excluded because of low eye-tracking data quality from the MRI session (> 3 standard deviations below the mean number of usable trials across participants). For one additional participant, one run of the search task due to excessive head motion.

The final sample of 32 participants consisted of 16 male individuals, 15 female individuals, and 1 non-binary individual. Sex/gender information was determined based on participant self-report. Sex/gender was not considered as part of the study design; evaluating sex/gender differences fell outside the primary scope and hypotheses of the study, thus no analyses of sex/gender differences were conducted. Participants included 11 individuals who identified as White, 9 who identified as Hispanic/Latino and White, 3 who identified as Black or African-American, 1 who identified as Hispanic/Latino and Black or African-American, 5 who identified as Asian, and 3 who identified as multiracial. Participants were 18–31 years old (median = 21) and had 12–20 years of education (median = 15.5). All participants were right-handed, had normal or corrected-to-normal visual acuity, normal color vision, and no MRI contraindications.

### Stimuli

#### Scenes

There were 24 critical scene stimuli, comprising 12 pairs of highly similar scene images (pairmates). Each pair belonged to a distinct outdoor scene category: apple orchards, arches, barns, canyons, castles, cities, gazebos, lighthouses, ponds, roller coasters, vineyards, and waterfalls. The two scene pairmates from each category were handpicked to have high levels of perceptual similarity. Scenes were sourced from the internet and included several of the categories used in Favila et al. 2016. We validated the perceptual similarity of our similar scenes by running the images through three convolutional neural network models (alexnet, vgg16, and resnet50) that approximate high-level visual cortex^84–86^ and were pretrained to classify scenes. We used late-layer activations from these models to compute scene similarity matrices for our stimulus set. Pairmate similarity distributions were higher than and non-overlapping with non-pairmate similarity distributions in all models, validating our pairmate similarity structure. Beyond these critical scenes, an additional 56 outdoor scenes were used as novel images in the fMRI task only. These scenes were hand-selected to have low perceptual similarity with the scene pair stimuli and with each other. Throughout the experiment, scenes were displayed in the center of the screen and were sized to be 15 x 15 degrees of visual angle. Participants never saw any two scenes presented simultaneously for comparison. Figures contain public domain images intended to be representative of the stimuli in the experiment. The actual stimuli used in the experiment can be downloaded with the data.

#### Locations

Eight isoeccentric (eccentricity = 3.3 degrees of visual angle) spatial locations were evenly sampled across polar angles (theta = 22.5, 67.5, 112.5, 157.5, 202.5, 247.5, 292.5, or 337.5 degrees). During learning, locations were always visualized as a white circle with a green outline centered at one of these locations and with a radius of 0.75 degrees of visual angle. Participants never saw the eight locations highlighted simultaneously.

#### Scene Distortions

Spatially localized distortions were created in the scene images by vertically displacing a local patch of pixels according to a sinusoidal function. The distortion was smoothly modulated by a fixed 2D Gaussian function so that the distortion was strongest at the center and faded gradually outwards. Perceptually, this resulted in a small circular patch of the image looking wavy with no hard boundary between the distorted patch and the rest of the image. For each of the 24 critical scene stimuli and each of the 56 novel scene stimuli, we created eight different distorted images. Each version of an image contained a single distorted location centered at one of the eight isoeccentric spatial locations used in the competitive learning task. To create each distorted image, we initially used a sinusoid with a fixed spatial frequency, a fixed amplitude, and a randomly selected phase, centered at the distortion location and smoothed with a 2D Gaussian. Then, to equate the difficulty of identifying distortions across different images and across different locations within an image, we ran a separate online experiment. Online participants viewed the initial images one at a time for 3 s each and were instructed to click on the distortion as quickly as possible. Based on participants’ ability to detect the distortions, we separately adjusted the amplitude of the sinusoid (distortion strength) up or down for each image and location, targeting an accuracy of 50% for each image and location across participants. We did this iteratively across four rounds of data collection, resulting in final distortion strengths that were tailored to each location within each image. We purposely chose a relatively modest accuracy threshold for this norming study to ensure that detecting the distortions was difficult enough to encourage participants to use memory to guide their attention in the main search task.

### Experimental Design and Procedure

The experiment consisted of two sessions, which occurred on consecutive days. In the first session (behavior only), participants performed a competitive learning task during which they acquired competing scene-location associations. The following day, participants performed a memory-guided visual search task while being scanned with fMRI. This task required participants to use the scene-location associations they acquired in the first session to improve their search performance. Participants also performed a population receptive field mapping task (see Session 2: Retinotopic Mapping Task (fMRI)) and localizer task (not analyzed here) while being scanned. All tasks were implemented in PsychoPy. Experimenters were not blinded to the design or hypotheses. Design and procedure details for each session and task are described below.

#### Session 1: Competitive Learning Task (Behavioral)

The goal of the first session was for participants to learn competing scene-location associations. There were three types of blocks in this session: scene exposure, study, and test. Participants’ eye movements were collected continuously in all three block types.

##### Scene Exposure

Participants began the session with two blocks of scene exposure. They were instructed that this was an opportunity to become familiar with the scenes before learning the scene-location associations, and that some scenes would be similar to each other. In each scene exposure block, participants were presented with each of the 24 critical scenes one time. Scenes were presented in a random order with the constraint that pairmates not be presented back to back. Each scene was presented on a gray background for 2 s. Participants’ eye movements were unrestricted during scene presentation. A 2 s central fixation interval occurred in between trials. Fixation on the central dot during the intertrial interval was enforced with a beep in response to fixation breaks.

##### Scene-Location Associations

After completing the scene exposure blocks, participants acquired 24 scene-location associations. Critically, some of these memories competed with each other because of the pairmate structure of the scenes. Scene-location associations were determined according to the following procedure. Each of the 24 critical scenes was assigned to predict one of the eight spatial locations. This was done randomly for each participant such that: 1) pairmate scenes were never associated with the same location; 2) the distance between two pairmates’ associated locations (45, 90, 135, or 180 degrees) was balanced; 3) all eight locations were sampled evenly. We refer to the location associated with a scene as that scene’s target location and the location associated with the pairmate scene as the competitor location.

##### Study and Test Blocks

Participants acquired the scene-location associations through study and test blocks. During study blocks, participants were presented with scene-location associations one time each and were instructed to explicitly encode the associations to the best of their ability. At the start of each study trial, a scene was presented on a gray background for 2 s, followed by that scene’s associated location for 2 s. The location (see Methods/Stimuli/Locations) was overlaid on a dark gray box the same size as the scene. Participants’ eye movements were unrestricted during scene and location presentation. A 2 s enforced central fixation interval occurred in between study trials. Study trial order was randomized within a block with the constraint that scene pairmates not be presented in consecutive trials. During test blocks, participants were tested on their memory for the scene-location associations one time each. They were instructed to recall the associations as accurately as possible and to guess if necessary. At the start of each test trial, a scene cue was presented on a gray background for 2 s, followed by a 2 s enforced central fixation interval. The response period began immediately after this with the onset of a blank dark gray box the same size as the scene. Participants were instructed to make a memory-guided saccade to the cued scene’s associated location during this period and to hold their gaze at the remembered location. The response period lasted 2 s and the participant’s average eye position in the final 200 ms of the response period (excluding blinks) was recorded as their response. Angular error was computed as the angular distance between this response and the center of the correct location. Immediately following the response period and regardless of the participant’s response, the correct location was presented for 1 s as feedback. Participants’ eye movements were unrestricted during scene presentation and location feedback. A 2 s enforced central fixation interval occurred in between test trials. Test trial order was randomized within a block with the constraint that scene pairmates not be presented in consecutive trials. At the end of each test block, participants were shown a block feedback screen that reported their average angular error as well as the percentage of trials for which their response was closer to the target location than to the competitor location.

##### Block Order and Performance Evaluation

To ease the difficulty of the task, participants first performed three consecutive study-test rounds on 12 scene-location associations (6 scene pairs). They then repeated the process for the remaining 12 scene-location associations. After performing the second set of three study-test rounds, participants performed two test blocks in which they were tested on all 24 scene-location associations. If a participant achieved an average angular error of 20 degrees or less across these two blocks, the learning task was terminated. If a participant did not meet this threshold, they continued performing test blocks until the average of their last two blocks met this threshold or until the end of the 1.5 hour session was reached. We targeted these extra test blocks toward poorly learned pairs by dropping well-learned pairs from the blocks. Well-learned pairs were pairs for which the participant responded within 20 degrees of angular error to both pairmates on two consecutive blocks.

#### Session 1: Distortion and Search Practice (behavioral)

At the end of Session 1, participants did two short practice tasks that prepared them for the scanned search task in the next session. Participants were told that their task the next day would involve searching for distortions hidden in scenes. They were then given a short task designed to introduce them to what these distortions looked like. Participants viewed 30 example images with representative distortions, one at a time. The task was self-paced, allowing participants to search for each distortion for as long as they wanted before choosing to reveal the location of the distortion. None of the distorted images in this task appeared in the next session. Then, participants were instructed on the search task (see next section) and practiced a modified version of it with the 24 critical scenes while their eye movements were recorded. In this modified version, participants were given longer to search for the distortions (2.75 s vs. 1.25 s in the fMRI task) and all trials were valid.

#### Session 2: Memory Refresher and Search Practice (behavioral)

At the start of session 2, immediately before entering the scanner, participants did two short practice tasks to prepare them for the scanned search task. First, participants were re-tested on the scene-location associations that they learned the previous day. They then practiced the version of the search task that they would perform in the scanner. Both of these tasks used an eye-tracking setup equivalent to the one used in session 1, but located in the same building as the MRI facility. The exact number of practice blocks/trials was based on participant performance and comfort with the task.

#### Session 2: Memory-guided Visual Search Task (fMRI)

The goal of the second session was for participants to use their memories of the scene-location associations to guide performance in a visual search task. Participants completed the search task while being scanned with functional MRI (see MRI Acquisition) and while their eye movements were recorded continuously (see Eye Tracking Data Acquisition).

##### Task Design

The purpose of the task was to measure participants’ ability to use competing memories to guide online attentional behavior. To encourage reliance on memory, we designed a visual search task that was challenging to perform under the chosen task parameters (high search difficulty and short stimulus presentation). On each trial of this task, participants had 1.25 s to search a scene image for a small local distortion (see Stimuli/Scene Distortions). Under these conditions and based on our distortion norming experiment and other preliminary pilot data, participants were expected to locate the distortion on fewer than half of trials. Critically, we embedded structure in the task that allowed participants to improve their search performance if they could recall the scene-location associations they learned the previous day. Specifically, consecutive trials were linked such that each scene was both a search array (i.e. it contained one distortion) *and* a cue for the subsequent trial. If participants saw one of the 24 critical scene stimuli from session 1 in the trial sequence, that meant that the scene’s associated target location was the most likely location of the distortion on the next trial. This manipulation encouraged participants to recall the prior scene’s target location during the intertrial interval to predict the distortion location on the upcoming trial. There were three conditions of interest: validly cued trials, invalidly cued trials, and no-prediction trials. On validly cued trials, which were 75% of all trials, the distortion was placed in the previous scene’s target location. On invalidly cued trials, which were 12.5% of all trials, the distortion was placed in the location associated with the previous scene’s competitor (i.e., the target location of the previous scene’s pairmate). On no-prediction trials, which were 12.5% of all trials, the distortion was placed randomly in one of the eight possible distortion locations. These trials were always preceded by one of the trial-unique novel scenes, which had no associated target location (see Stimuli/Scenes). Trial orders were constructed such that the critical manipulation between scene identity and the next trial’s distortion location was the only exploitable structure. Scenes only predicted distortions on the *upcoming* trial; there was no link between scene identity and the location of the distortion on that particular trial. With the exception that scene pairmates were never presented in back-to-back trials, upcoming scene identity was not predictable. All of the 24 critical scene stimuli were equally likely to be validly cued and (within a tolerance) to be valid cues. All eight locations were equally likely to be the learned target location relevant for the next trial. Distortions occurred at all eight locations with approximately equal probability.

##### Task Parameters and Procedure

Participants performed eight consecutive runs of the search task in the MRI scanner with short breaks in between each run. On each trial of the task, a scene was presented on a gray background for 1.25 s. Every scene contained one local distortion, which occurred at one of eight isoeccentric locations (see Stimuli/Scene Distortions). The onset of scene presentation cued participants to begin their search. Participants were instructed to move their eyes to find each scene’s distortion as quickly as possible and to fixate the distortion when they found it. Participants’ entire eye movement trajectory was recorded, but for the purpose of generating feedback, the final 200 ms of their eye position was recorded and averaged excluding blinks. If this position was within 20 degrees of angular error from the center of the distortion location, the trial was considered correct. At the offset of scene presentation, an intertrial fixation interval began. This interval was randomly selected to be 4.75 s, 6.25 s, or 7.75 s with equal probability. For the first .75 s of the interval, the central fixation dot was green (if the previous trial was correct) or red (if the previous trial was incorrect), and for the rest of the interval it was white. Participants were explicitly instructed of the contingency between consecutive trials and were told to aim for 70% accuracy by relying on their memories to predict the location of upcoming distortions. They were also instructed to maintain fixation on the central fixation dot while preparing for the next trial. At the end of each run, participants were shown a feedback screen that reported their average accuracy for that run. Each run took a little over seven minutes to complete and contained 56 trials. The first trial in each run was discarded from behavioral analyses since it was not cued. Among the 55 cued trials per run, there were 41 validly cued trials, 7 invalidly cued trials, and 7 no-prediction trials. The 55 cued trials always consisted of two presentations each of the 24 critical scenes and one presentation each of 7 trial-unique novel scenes. Across all eight runs of the task, each of the 24 critical scenes was presented 16 times; 12 of these presentations were validly cued trials, 2 were invalidly cued trials, and 2 were no-prediction trials.

#### Session 2: Retinotopic Mapping Task (fMRI)

After completing the search task, each participant completed four identical retinotopic mapping runs. Stimuli and task parameters for the retinotopic mapping session were based on those reported previously^87,88,42^, but a smaller portion of the visual field was mapped. During each functional run, bar apertures on a uniform gray background swept across the central 15 degrees of the participant’s visual field (a circular aperture with a radius of 7.5 degrees of visual angle). Each sweep began at one of eight equally spaced positions around the edge of the circular aperture, with the bar oriented perpendicularly to the direction of the sweep. During horizontal and vertical sweeps, the bar moved from one side of the aperture to the other at a constant rate. During diagonal sweeps, the bar moved at a constant rate until it reached the midpoint of the aperture; it then disappeared and was followed by a blank period that allowed for measurement of baseline neural activity. The bar position was incrementally updated every 1 s, such that a full-field sweep across the entire aperture took 24 s. Half-field sweeps took 12 s and subsequent blank periods took 12 s (24 s total). One functional run contained 8 back-to-back sweeps, taking 192 s in total. Bar apertures were a constant width and contained a grayscale pink noise background with randomly placed faces, scenes, objects, and words at a variety of sizes. Noise background and stimuli were updated at a frequency of 3Hz. Each run of the task had identical timing, bar positions, sweep order, and stimuli. Participants were instructed to maintain fixation on a central dot and to use a button box to report whenever the dot changed color from blue to green or green to blue. Color changes occurred 24 times per run, and their timing was randomized for each participant and run.

### Eye Tracking Data Acquisition

Eye tracking data were collected using an Eyelink 1000 Plus infrared video-based system by SR Research. For all sessions, the camera tracked the participant’s right eye at 1000 Hz in a head-stabilized position. During session 1, participants were stabilized in a chinrest and the camera was positioned below the task display. During session 2, participants were stabilized in the head coil and an MRI-compatible long-range enabled Eyelink 1000 Plus system was positioned outside the scanner bore and beneath the MRI display such that the participant’s eye could be tracked through the head coil-mounted mirror. In both sessions we calibrated the eye tracker using a 9 point calibration scheme at the start of each task block or scanner run, as needed. Re-calibration was performed when the experimenters observed high levels of noise in tracked gaze position during fixation periods, when gaze tracking was lost intermittently during the search interval, or when validation tests failed.

### Eye Tracking Preprocessing

Raw gaze data from session 1 and session 2 were preprocessed to identify saccades during task periods of interest. For session 1, we extracted gaze data from the response period of test trials and converted the gaze coordinates from pixels into degrees of visual angle. We removed the 200 ms prior to blinks and the 200 ms after blinks to remove artifacts. We used smoothed gaze data to define saccades as eye movements with a velocity of at least 30 degrees of visual angle/s and an amplitude of at least 0.25 degrees of visual angle^89^. Fixations were defined to be periods of stable gaze position occurring between saccades, excluding blinks. We corrected saccade trajectories based on the longest fixation of the pretrial central fixation period to account for drift. For our analyses, we dropped all saccades that landed within 1 degree of visual angle from central fixation and analyzed the remaining saccades (see Methods/Behavioral Analyses for more details). For session 2, we performed the same procedure, but on gaze data from the scene presentation period of search trials.

To assess the reliability of eye-tracking data collected in the scanner, we computed several data quality measures on the preprocessed gaze estimates from the search task. Calibration was performed on 43% of runs, with wide variability between participants (min = 12.5%, max = 100%), depending on experimenter assessment of eye-tracking quality and the results of validation tests at the start of every run. The mean final calibration error (the average deviation between the measured and intended target positions during the final calibration) was 0.47 degrees of visual angle, with an average run-to-run SD of 0.27 degrees of visual angle. After preprocessing (including blink removal), data completeness during the critical search period was high across participants (mean valid samples = 88.6%, SD = 6.1%). Because gaze position changed rapidly during the search period, we quantified spatial precision from the ITI fixation period, when participants maintained a stable gaze. For each run, the standard deviation of x- and y-coordinate gaze samples was computed, and spatial precision was defined as the mean of these two values. Spatial precision during this period was 0.46 degrees of visual angle (SD = 0.18 degrees of visual angle), indicating low sample-to-sample variability and good measurement quality.

For additional checks of our eye tracking data, we quantified all saccades that participants initiated during both scene presentation (when eye movements were required) and the ITI (when participants were supposed to maintain fixation). Participants typically made 3 saccades per search trial (median across all runs and all participants). They made significantly fewer saccades on trials in which they found the distortion than trials in which they didn’t (t(31) = −6.17; p < 0.001; d_z_ = 0.94; successful - unsuccessful = −0.65; 95% CI = [−0.87, −0.44]), indicating that they understood the task objective and stopped searching when they located the distortion. Participants typically broke fixation during the ITI on only 3.5 of out 56 trials per run (median across all runs and all participants), indicating that they understood when to withhold and make eye movements. The pattern of results for all ITI analyses of brain activity was unaffected when trials associated with broken fixation during the ITI were excluded from analysis.

### MRI Acquisition

All MRI data were acquired on a 3T Siemens Magnetom Prisma system located at Columbia University’s Zuckerman Mind Brain Behavior Institute. Functional images were acquired using a Siemens 64-channel head/neck coil and a multiband EPI sequence (repetition time = 1.5s, echo time = 30ms, flip angle = 65°, acceleration factor = 3, voxel size = 2 x 2 x 2mm, phase encoding direction = posterior to anterior). Sixty-nine oblique axial slices (14° transverse to coronal) were collected in an interleaved order. Each participant also underwent a T1-weighted anatomical scan (resolution = 1 x 1 x 1mm) using a magnetization-prepared rapid acquisition gradient-echo (MPRAGE) sequence. Double-echo gradient echo images were acquired to estimate the susceptibility-induced distortion present in the functional EPIs. In two participants, an opposite phase encoded field map sequence (anterior to posterior and posterior to anterior) was collected instead.

### MRI Preprocessing

Results included in this manuscript come from preprocessing performed using fMRIPrep 21.0.0^90,91^ (RRID:SCR_016216), which is based on Nipype 1.6.1^92^ (RRID:SCR_002502).

#### Preprocessing of B0 Inhomogeneity Mappings

A B_0_ nonuniformity map (or fieldmap) was estimated from the phase-drift map(s) measure with two consecutive GRE (gradient-recalled echo) acquisitions. The corresponding phase-map(s) were phase-unwrapped with prelude (FSL 6.0.5.1:57b01774). Due to a protocol change, for two participants, fieldmaps were estimated based on two (or more) echo-planar imaging (EPI) references.

#### Anatomical Data Preprocessing

The T1-weighted (T1w) image was corrected for intensity non-uniformity (INU) with N4BiasFieldCorrection^93^, distributed with ANTs 2.3.3^94^ (RRID:SCR_004757), and used as T1w-reference throughout the workflow. The T1w-reference was then skull-stripped with a Nipype implementation of the antsBrainExtraction.sh workflow (from ANTs), using OASIS30ANTs as target template. Brain tissue segmentation of cerebrospinal fluid (CSF), white-matter (WM) and gray-matter (GM) was performed on the brain-extracted T1w using fast (FSL 6.0.5.1:57b01774, RRID:SCR_002823^95^). Brain surfaces were reconstructed using recon-all (FreeSurfer 6.0.1, RRID:SCR_001847^96^), and the brain mask estimated previously was refined with a custom variation of the method to reconcile ANTs-derived and FreeSurfer-derived segmentations of the cortical gray-matter of Mindboggle (RRID:SCR_002438^97^). Volume-based spatial normalization to one standard space (MNI152NLin2009cAsym) was performed through nonlinear registration with antsRegistration (ANTs 2.3.3), using brain-extracted versions of both T1w reference and the T1w template. The following template was selected for spatial normalization: ICBM 152 Nonlinear Asymmetrical template version 2009c^98^ (RRID:SCR_008796; TemplateFlow ID: MNI152NLin2009cAsym).

#### Functional Data Preprocessing

For each of the BOLD runs per participant, the following preprocessing was performed. First, a reference volume and its skull-stripped version were generated using a custom methodology of fMRIPrep. Head-motion parameters with respect to the BOLD reference (transformation matrices, and six corresponding rotation and translation parameters) are estimated before any spatiotemporal filtering using mcflirt (FSL 6.0.5.1:57b01774^99^). The estimated fieldmap was then aligned with rigid-registration to the target EPI (echo-planar imaging) reference run. The field coefficients were mapped on to the reference EPI using the transform. BOLD runs were slice-time corrected to 0s for retinotopic mapping runs and 0.708s for memory-guided search runs using 3dTshift from AFNI^100^ (RRID:SCR_005927). The BOLD reference was then co-registered to the T1w reference using bbregister (FreeSurfer) which implements boundary-based registration^101^. Co-registration was configured with six degrees of freedom.

Several confounding time series were calculated based on the preprocessed BOLD: framewise displacement (FD), DVARS and three region-wise global signals. FD was computed using two formulations following Power (absolute sum of relative motions, Power et al., 2014) and Jenkinson (relative root mean square displacement between affines^99^. FD and DVARS are calculated for each functional run, both using their implementations in Nipype (following the definitions by Power et al.^102^. The three global signals are extracted within the CSF, the WM, and the whole-brain masks. Additionally, a set of physiological regressors were extracted to allow for component-based noise correction (CompCor^103^). Principal components are estimated after high-pass filtering the preprocessed BOLD time series (using a discrete cosine filter with 128s cut-off) for the two CompCor variants: temporal (tCompCor) and anatomical (aCompCor). tCompCor components are then calculated from the top 2% variable voxels within the brain mask. For aCompCor, three probabilistic masks (CSF, WM and combined CSF+WM) are generated in anatomical space. The implementation differs from that of Behzadi et al. in that instead of eroding the masks by 2 pixels on BOLD space, the aCompCor masks are subtracted a mask of pixels that likely contain a volume fraction of GM. This mask is obtained by dilating a GM mask extracted from the FreeSurfer’s aseg segmentation, and it ensures components are not extracted from voxels containing a minimal fraction of GM. Finally, these masks are resampled into BOLD space and binarized by thresholding at 0.99 (as in the original implementation). Components are also calculated separately within the WM and CSF masks. For each CompCor decomposition, the *k* components with the largest singular values are retained, such that the retained components’ time series are sufficient to explain 50 percent of variance across the nuisance mask (CSF, WM, combined, or temporal). The remaining components are dropped from consideration. The head-motion estimates calculated in the correction step were also placed within the corresponding confounds file. The confound time series derived from head motion estimates and global signals were expanded with the inclusion of temporal derivatives and quadratic terms for each^104^. Frames that exceeded a threshold of 0.5 mm FD or 1.5 standardized DVARS were annotated as motion outliers.

The BOLD time series were resampled into standard space, generating a preprocessed BOLD run in MNI152NLin2009cAsym space. The BOLD time series were resampled onto the following surfaces (FreeSurfer reconstruction nomenclature): fsaverage, fsnative. All resamplings can be performed with a single interpolation step by composing all the pertinent transformations (i.e. head-motion transform matrices, susceptibility distortion correction when available, and co-registrations to anatomical and output spaces). Gridded (volumetric) resamplings were performed using antsApplyTransforms (ANTs), configured with Lanczos interpolation to minimize the smoothing effects of other kernels^105^. Non-gridded (surface) resamplings were performed using mri_vol2surf (FreeSurfer).

Many internal operations of fMRIPrep use Nilearn 0.8.1^106^, RRID:SCR_001362), mostly within the functional processing workflow. Note that the analyses presented in this paper only make use of preprocessed BOLD data in native T1w and fsnative spaces.

#### Copyright waiver

The above boilerplate text was automatically generated by fMRIPrep with the express intention that users should copy and paste this text into their manuscripts unchanged. It is released under the CC0 license.

### pRF Model Fitting

pRF model fitting was conducted using vistasoft using an approach described by Dumoulin & Wandell^107^. We took the preprocessed BOLD time series from the retinotopic mapping task in native surface space and upsampled them to a temporal resolution of 1.0s to match the rate at which the bar stimulus moved. We then averaged them across runs. Using this average time series, we estimated the best fitting pRF for each vertex. Each pRF was modeled as a circular 2D Gaussian, parameterized by values *x, y*, and *σ*. The *x* and *y* parameters specify the center position of the 2D Gaussian in the visual field. These parameters can be re-expressed in polar coordinates: *r* (eccentricity), *θ* (polar angle). The *σ* parameter, the standard deviation of the 2D Gaussian, specifies the size of the receptive field. Stimulus images from the retinotopic mapping task (Methods/Experimental Design and Procedure/Session 2: Retinotopic Mapping Task) were converted to contrast apertures. Candidate pRFs were multiplied pointwise by the stimulus apertures to yield a predicted time course of activation by the stimulus, which was then convolved with a double-gamma hemodynamic response function to predict the BOLD response in each vertex. Using an iterative coarse-to-fine fitting approach, we estimated the *x, y*, and *σ* parameters for each vertex that, when applied to the drifting bar stimulus, minimized the difference between the observed and predicted BOLD time series.

### Regions of Interest

The hippocampus was defined bilaterally in each participant’s native anatomical space using Freesurfer. For some analyses, we divided this ROI into anterior and posterior components. We did this by taking a hippocampal ROI in MNI space and dividing it into anterior and posterior components at y = −21, following the convention in Poppenk et al.^38^. We then reverse-normalized this ROI into native T1 space and used the transformed anterior-posterior boundary in native space to segment each participant’s whole hippocampal Freesurfer ROI. We used this approach to ensure that our anterior and posterior hippocampal ROI summed to our original whole hippocampus ROI.

Visual cortex regions of interest were defined by visualizing the pRF parameters on each participant’s native cortical surface and hand-drawing boundaries between visual field maps. For each of V1, V2, and V3, we drew boundaries at polar angle reversals, following established practice^108,109^. We combined bilateral V1, V2, and V3 surface masks into a single early visual cortex (EVC) ROI. This ROI was projected to native T1 space to create a volumetric EVC ROI for some analyses.

We additionally defined several regions implicated in scene-processing to serve as control ROIs: PPA, RSC, and PHC. To define PPA, we selected the CoS-places mask from a published functional group atlas^110^ and registered it to each participant’s native surface space. We then projected this surface-based ROI to native T1 space to create a volumetric ROI. To define RSC, we relied on a neurosynth contrast previously used in one of our papers^111^. We defined PHC anatomically, according to agreed upon protocols for segmenting the medial temporal lobe (see Aly & Turk-Browne^20^). Both RSC and PHC were defined in MNI space and then reverse-normalized into native T1 space for each participant. Finally, we defined the Dorsal Attention Network (DAN) to serve as a control ROI. To do this, we transformed the group-average 7 Network parcellation from Yeo et al.^112^ into native surface space and then native volumetric space and selected network 3.

### Behavioral Analyses

We analyzed the end points of the first and last saccades executed during task periods of interest (see Methods/ Eye Tracking Data Acquisition and Methods/Eye Tracking Preprocessing for details about how gaze positions were estimated). Saccade end points were analyzed with respect to each trial’s spatial locations of interest: the target location, the competitor location, and the control location. Target and competitor locations were defined based on associative learning in session 1. The control location was defined to be the same distance away from the target location as the competitor location, but in the opposite direction. Thus for target and competitor locations that were 180 degrees away, the control location was undefined. We computed several metrics with respect to these locations. First, we assigned each saccade to the target, competitor, or control locations if it was closer to that location than to any of the other studied locations (i.e. if the angular error between the saccade end point and the center of the relevant location was less than 22.5 degrees). We assessed the probability of these response types across conditions of interest for both the associative learning task and the search task. For the search task, we also analyzed the angular error between the saccade end point and the center of the distortion location as a continuous measure. Finally, for the search task, we also considered saccade accuracy. A saccade was considered accurate (alternatively referred to as a search success) if it fell within 22.5 degrees of the distortion location. A saccade was considered an interference error if it fell within 22.5 degrees of the competitor location. Finally, a saccade was considered an other error if it fell in any of the locations other than the target or competitor location.

### Pattern Similarity Analyses

To estimate the pattern of activity evoked by similar scenes, we conducted a voxelwise GLM analysis of each participant’s preprocessed time series data from the search task in native T1 space. Each model contained a regressor of interest for each stimulus in the experiment (24 critical scenes [12 pairs of similar scenes] and 56 novel scenes). Events within these regressors were constructed as boxcars with a duration of 1.25 s (scene presentation duration), convolved with a double-gamma hemodynamic response function. Six realignment parameters and cosine drift parameters were included as nuisance regressors to control for motion confounds and signal drift. No spatial smoothing was performed. We included all eight runs of the memory-guided search task in each GLM, representing 16 presentations of each of the 24 critical scenes, with the exception of one participant. One run was excluded from this participant because >10% of TRs had a framewise displacement of > 1mm. First level models were estimated using nilearn^106^ and contrasts for each of the 24 critical scenes versus baseline were computed. While regressors for the novel scenes were included in these models, these parameter estimates were not analyzed further. This procedure yielded t-maps representing the activation elicited by viewing each of the 24 critical scenes (12 pairs of similar scenes) for each participant.

To compute scene pair pattern similarity, we calculated the Fisher z-transformed Pearson correlation between the t-maps for all unique pairwise combinations of the 24 scenes. For each stimulus category (e.g., arches), we averaged all the correlations between nonpairmate scenes that included a scene from that category (e.g., arch 1 and barn 1; arch 2 and barn 1, etc). We subtracted this value from the correlation of the two pairmate scenes from that category (e.g., arch 1 and arch 2). Using this metric, lower values indicate that scene pairmates are represented more distinctly compared to other scenes and higher values indicate that scene pairmates are represented more similarly. To assess the overall tendency of a brain region to differentiate similar scenes, we conducted these analyses separately within brain regions of interest (hippocampus plus control regions) and averaged the scene pair pattern similarity values across the 12 scene pairs to produce one value per participant. To assess the relationship between scene separability and variables of interest, we sorted the scene pairs by their scene pair pattern similarity value in a particular ROI and divided them into more and less distinct halves. For some analyses, we considered scene pair similarity continuously; in this case, we normalized scene pair similarity values for an ROI within participants.

In addition to our ROI-based analyses, we also conducted an exploratory whole-brain searchlight analysis that assessed the relationship between scene pair pattern similarity and behavior across the entire brain. For each participant, we computed our scene pair pattern similarity score in searchlights of radius 2 voxels, covering the entire brain in native T1 space. For each searchlight, we sorted scene pairs according to their scene pair similarity value into more and less distinct halves. We then calculated a paired t-statistic describing the difference in search error between these scene groups (equivalent to the analysis reported in Figure 3C, but conducted within searchlights). We transformed these native space t-maps into MNI space and spatially smoothed them with a 4mm Gaussian kernel. We then ran a permutation test using FSL’s *randomise* function (1000 permutations) on t-values falling within gray matter to produce uncorrected threshold-free cluster enhancement p values.

### Visual Cortex Preparatory Responses

We examined preparatory BOLD responses during the inter-trial interval (ITI) period of the search task by first removing the responses evoked by the preceding scene presentation and then segregating residual ITI responses according to their visual field position. To remove the visually-evoked responses, we conducted a voxelwise GLM analysis of each participant’s preprocessed time series data from the search task in native surface space. (Note that we use the term voxel in the Results for convenience to readers, but pRF analyses are done on vertices in surface space rather than voxels in volume space). Each model contained two regressors of interest: one for presentations of the 24 critical scenes (12 pairs of similar scenes) and one for presentations of the novel scenes. These regressors and nuisance regressors were constructed using the same procedure as described in the Pattern Similarity Analyses subsection above, and the models were estimated using the same data inclusion criteria. We took the residual time series after estimating this model and z-scored each vertex across time and then each time point across vertices.

Using these residual time series, we isolated the time points corresponding to the first 3 TRs of the blank ITI of the search task after accounting for hemodynamic lag. This corresponded to TRs 4, 5, and 6 from the prior scene onset. We chose 3 TRs because all trials had at minimum 3 TRs for the ITI; this allowed us to keep the amount of data constant across trials that varied in ITI duration. We averaged the residual time series across these TRs for each trial, generating one ITI BOLD response value per vertex per trial. We considered only valid trials for further analyses. To understand how these ITI responses corresponded to visual space, we organized these responses according to the values of the parameter estimates derived from the pRF model. We first selected the subset of EVC vertices that were potentially responsive to the 8 target locations in our experiment. We included vertices whose pRFs met 3 criteria: (1) they had eccentricity values between 0.5 and 10 degrees, indicating they were responsive to the display screen excluding central fixation; (2) the center of the 2D Gaussian was within 2*σ* of the circle along which target locations were aligned, indicating they were additionally responsive to target and competitor locations; and (3) their pRF model *R*^2^ was at least 0.1, indicating they exhibited spatial selectivity.

To visualize the ITI BOLD response of vertices with different pRFs^42^, we divided the selected vertices into 8 bins according to the polar angle distance between their pRF and the target memory location on the upcoming trial. We took each participant’s median ITI response within a bin before averaging across participants. This procedure produced a spatial response function that visualizes how strong the ITI response was in vertices coding for the upcoming target location and how quickly this response falls off over space. Note that for visualization purposes only, we visualized the same data at +180 and −180 degrees away, so that the target location appears centered. Visualizations examining competitor location responses during the ITI followed the same procedure but binned vertices according to their distance from the competitor location. As an alternative visualization, we also assigned each of the 8 angular distance bins one of the following labels, separately for each participant and trial: target, competitor, control, or other. We then averaged the BOLD response separately for each of the four labels.

### Statistics

We evaluated the statistical reliability of our behavioral and pattern similarity results using paired t-tests, repeated measures ANOVAs, and Pearson correlations. We report 95% CIs on the difference in means (for t statistics) or for the parameter of interest (for Pearson correlations). We additionally report Cohen’s *d_z_* effect size for t statistics and partial eta-squared effect size for repeated measures ANOVAs. Reported p-values are two-tailed and evaluated at alpha = 0.05.

We used linear mixed effects models to assess the link between preparatory responses in visual cortex and 1) behavior; 2) hippocampal distinctiveness. For all models described below, we initially fit a model including the maximal random effects structure ^113^. However, in cases of non-convergence or singular fits, which indicate that the random effect structure is overparameterized for the complexity of the data, we removed random slopes until convergence was achieved. We always included a random intercept for each participant. We therefore report results from the most parsimonious model that converged and yielded stable parameter estimates^114^. We note that all significant effects identified in parsimonious (random intercept-only) models remain significant and consistent in direction when compared against the maximal models. Following best practices to reduce Type 1 error rate, we fit all models with Restricted Maximum Likelihood and used the Satterthwaite approximation to calculate degrees of freedom and report p values^115^. All models were implemented in the lme4^116^, lmerTest^117^, and emmeans^118^ packages for R.

To examine the relationship between preparatory responses in visual cortex and behavior on the subsequent trial, we fit linear mixed effects models to single trial data. For each valid trial and participant, the ITI BOLD response was extracted from the eight pRF bins described above and entered as the dependent variable. Trials in which no eye movement response was made on the subsequent trial were excluded from analysis. Behavioral outcome on the subsequent trial (success, interference, or other) and pRF location labels (target, competitor, control, or other) were included as fixed effects, along with their interaction. The most parsimonious model had the following specification: lmer(bold ∼ pRF_cond * behav_outcome + (1 | subj), data=valid_trials, REML=T). The pRF location x behavioral outcome interaction was first assessed using Type III F-tests. To probe the interaction, estimated marginal means were used to perform planned pairwise comparisons between pRF locations within each behavioral outcome. For each comparison, we report the estimated coefficient (β), 95% CI, t statistic, degrees of freedom, and p value.

To examine whether hippocampal memory representations predicted preparatory activity in visual cortex for 45 degree pairs, we fit a linear mixed-effects model relating hippocampal distinctiveness to ITI responses in different pRF voxel groups. As in the prior analysis, the ITI BOLD response extracted from each of the eight pRF bins for each valid trial and participant served as the dependent variable. Hippocampal distinctiveness was computed as pattern similarity at the level of each scene pair (see Methods/Pattern Similarity Analyses) and sorted in low distinctiveness (high pattern similarity) and high distinctiveness (low pattern similarity) groups. Hippocampal distinctiveness (low, high) and pRF location labels (target, competitor, control, or other) were included as fixed effects, along with their interaction. The most parsimonious model had the following specification: lmer(bold ∼ pRF_cond * sep_bin + (1 | subj), data=pairs45_data, REML=T). As before, the pRF location x hippocampal distinctiveness interaction was first assessed using Type III F-tests from the mixed effects model. To probe the interaction, estimated marginal means were used to perform planned pairwise comparisons between pRF locations within each hippocampal distinctiveness group. For each comparison, we report the estimated coefficient (*β*), 95% CI, t statistic, degrees of freedom, and p value.

To examine whether these effects generalized across pair distance, we fit additional models that tested 90 degree, 135 degree, and 180 degree pairs using the same fixed effects model structure. We also tested the three-way interaction between hippocampal distinctiveness, pRF location, and target-competitor distance. We used a similar mixed-effects modeling approach described above; however, for these analyses, control pRFs were omitted because some pair distances (e.g., 180°) did not allow for a valid definition of a control location, and including them would have resulted in missing cells and non-estimable parameters in the F test. This had no bearing on our primary effect of interest, which was in the relative activity of voxels coding for target vs. competitor locations. In addition, to assess the anatomical specificity of the distinctiveness effect, we fit additional models using data from ROIs besides the hippocampus. Finally, we examined univariate activity instead of our pattern similarity-derived distinctiveness measure. We tested the interaction between pRF location and univariate activity in the DAN using the same modeling framework. Univariate activity was estimated from the same GLM used for the pattern similarity analyses (see Methods/Pattern Similarity Analyses), with t statistics for each scene averaged over voxels in the DAN to provide a measure of univariate activity for each scene pair.

## Data Availability

Raw data have been deposited on OpenNeuro at https://doi.org/10.18112/openneuro.ds008366.v1.0.0119. Intermediate data needed to regenerate the figures is available on the Open Science Framework (OSF) at https://doi.org/10.17605/OSF.IO/C948A120.

## Code Availability

The code used to produce the analyses in this study is available on OSF at Acknowledgements

We thank Shelton Brister and Kaylee Wang for their assistance with data collection and the Alyssano Group and Brice Kuhl for their helpful feedback on this project.

## Author Contributions Statement

Conceptualization, S.E.F. and M.A.; Methodology, S.E.F. and M.A.; Software, S.E.F.; Investigation, S.E.F.; Formal Analysis, S.E.F.; Writing – Original Draft, S.E.F. and M.A.; Writing – Review & Editing, S.E.F. and M.A.; Funding Acquisition, M.A.; Resources, M.A.; Supervision, M.A.

## Supplementary Information

**Supplementary Figure 1.**
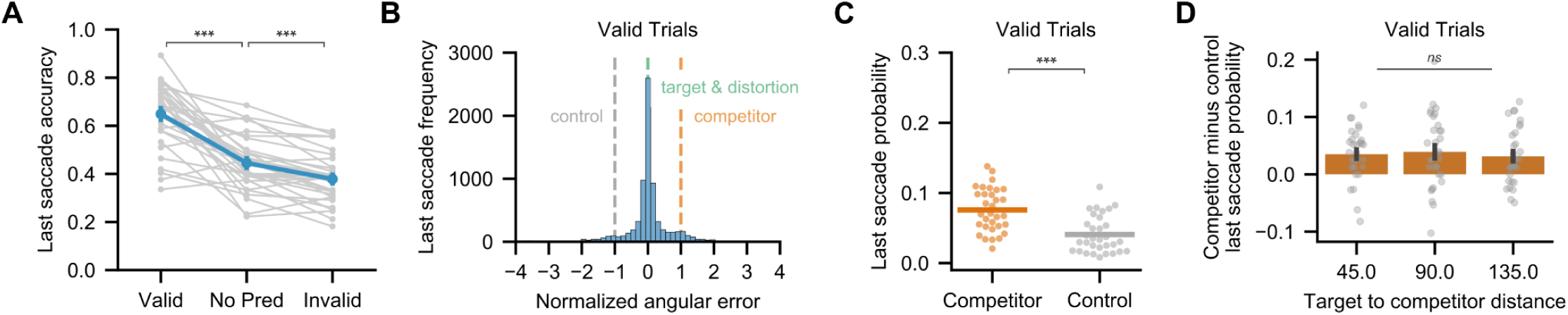
Last saccade behavioral data from memory-guided visual search task (session 2). **(A)** Relative to no-prediction trials, participants’ last saccade was more accurate on valid trials (t(31) = 7.34, p < 0.001, d_z_ = 1.62, valid - no-prediction = 0.20, 95% CI = [0.15, 0.26]) and less accurate on invalid trials (t(31) = −5.47, p < 0.001, d_z_ = 0.62, invalid - no-prediction = −0.067, 95% CI = [−0.09, −0.04]), demonstrating memory-guided search (N = 32 participants, two-tailed paired t-tests). Gray lines indicate individual participants; blue lines indicate across-participant mean ± SEM. **(B)** Last saccade angular error distribution on valid trials. Errors are normalized by target-competitor distance (x=0: target/distortion; x=1: competitor; x=−1: equidistant control). Saccades were most frequent near the target, but the distribution is asymmetric toward the competitor. **(C)** Last saccade probabilities for competitor and control locations from (B). Saccades to the competitor were more frequent than to the control (t(31) = 5.05, p < 0.001, d_z_ = 1.22, competitor - control = 0.035; 95% CI = [0.02, 0.05]; N = 32 participants, two-tailed paired t-test), suggesting that competition persisted into the search task. Dots indicate individual participants; lines indicate across-participant means. **(D)** Target-competitor distance did not significantly modulate last saccade likelihood to the competitor relative to control (F(2,62) = 0.265, p = 0.768, η^2^_P = 0.008, N = =32 participanths;_ repeated measures ANOVA). Bars indicate across-participant mean ± SEM. ns = p > 0.05; *** p < 0.001

**Supplementary Figure 2.**
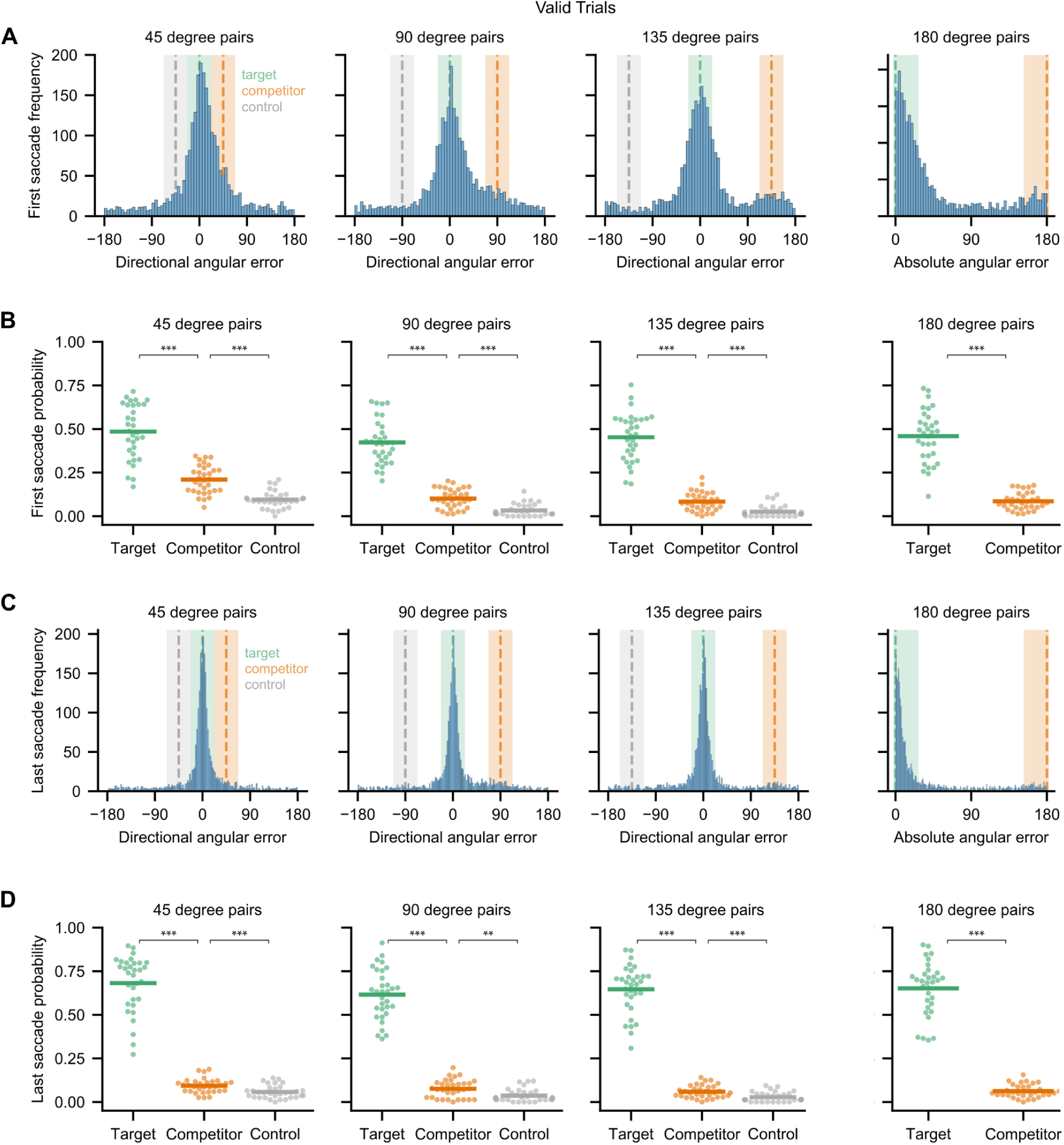
Behavioral data from memory-guided visual search task (session 2) separated by target-competitor distance. **(A)** The distribution of first saccade angular errors on valid trials, separately for each target-competitor distance. Zero degrees of error indicates a saccade made directly towards the target. For target-competitor distances of 45 degrees, 90 degrees, and 135 degrees, angular error is directional, such that positive values are toward the competitor and negative values are in the opposite direction toward a control location an equal distance away from the target as the competitor. For 180 degree pairs, no control location can be defined, so angular errors are collapsed across direction. Highlighted green areas represent the saccades that are identified to be at the target location. Orange and gray highlighted areas represent the saccades that are identified to be at the competitor and control locations, respectively. See Figure 2D for data normalized such that all target and competitor distances can be visualized at the same time. **(B)** Data from the target, competitor, and control shaded zones in (A) are re-plotted as probabilities for each participant. For every target-competitor distance, first saccades are made more often to the target than the competitor location. For the distances for which a control location can be defined, first saccades are made more often to the competitor than the control location. Dots represent individual participants and lines represent the across-participant mean. **(C)** Same as (A) but for the last saccade. **(D)** Same as (B) but for the last saccade. ** p < 0.01; *** p < 0.001

**Supplementary Figure 3.**
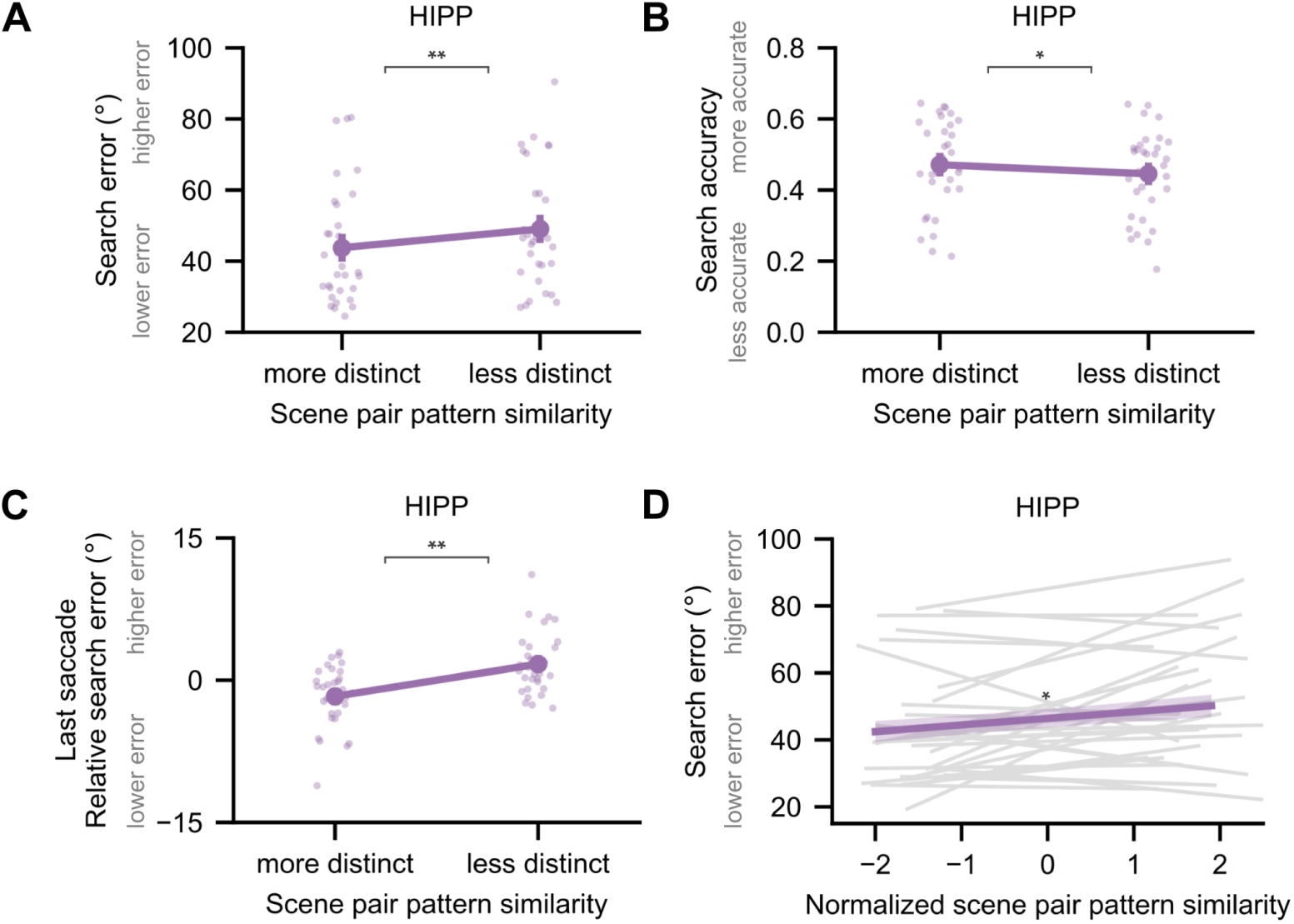
Distinct hippocampal representations of competing scenes predict multiple measures of memory-guided eye movements. **(A)** The relationship between pattern distinctiveness in the hippocampus (HIPP) and search error on the next trial (t(31) = −3.46, p = 0.002, d_z_ = 0.32, more distinct - less distinct = −5.3 degrees, 95% CI = [−8.43, −2.18]; N = 32 participants, two-tailed paired t-test), visualized without centering data within participants. See Figure 3C for the same search error values centered within participants. **(B)** The relationship between pattern distinctiveness in the hippocampus and behavior was robust to binarizing search error into search accuracy (t(31) = 2.04, p = 0.049, d_z_ = 0.21, more distinct - less distinct = 0.026, 95% CI = [0.0, 0.05]; N = 32 participants, two-tailed paired t-test). Search accuracy was defined as the proportion of trials with first saccade errors less than 22.5 degrees. **(C)** The relationship between pattern distinctiveness in the hippocampus and behavior was robust when considering search error on the last saccade rather than the first saccade (t(31) = −3.01, p = 0.005, d_z_ = 0.23, more distinct - less distinct = −3.4 degrees, 95% CI = [−5.73, −1.10]; N = 32 participants, two-tailed paired t-test). For panels A-C, small dots represent individual participants. Large points represent the across-participant mean ± SEM. **(D)** The relationship between pattern distinctiveness in the hippocampus and behavior was robust when considering pattern distinctiveness as a continuous rather than binary measure (t(315) = 2.38, p = 0.018, β = 1.98, 95% CI = [0.34, 3.62]; N = 384 scene pairs across 32 participants; two-tailed post-hoc contrast on a linear mixed effects model). The purple line and shading represent the slope and standard error of the fixed effect from a model predicting search error from scene pair pattern similarity (normalized within participants). Gray lines represent the best fitting line to each participant’s data. * p < 0.05; ** p < 0.01

**Supplementary Figure 4.**
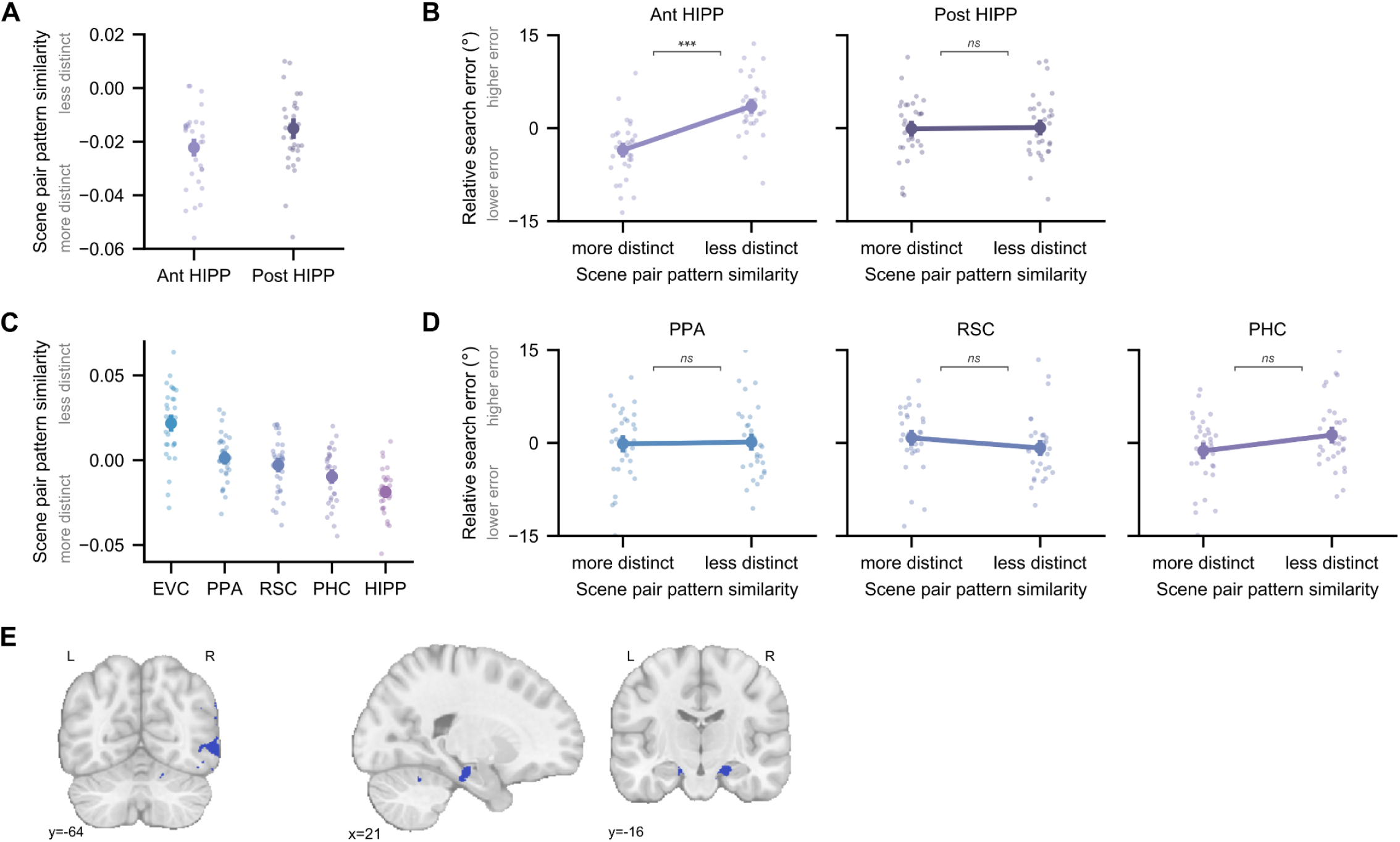
Relationship between representations of competing scenes and memory-guided eye movements in other brain regions. **(A)** Average scene pair pattern similarity in anterior hippocampus (Ant HIPP) and posterior hippocampus (Post HIPP). **(B)** Scene pair pattern similarity in the anterior hippocampus, but not posterior hippocampus, predicted lower error saccades (anterior hippocampus: t(31) = −4.33, p < 0.001, d_z_ = 0.43, more distinct - less distinct = −7.1 degrees, 95% CI = [−10.44, −3.75]; posterior hippocampus: t(31) = −0.11, p = 0.915, d_z_ = 0.01, more distinct - less distinct = −0.19 degrees, 95% CI = [−3.84, 3.45]; N = 32 participants, two-tailed paired t-tests). **(C)** Average scene pair pattern similarity in early visual cortex (EVC) and hippocampus (as reported in the main text), parahippocampal place area (PPA), retrosplenial cortex (RSC), and parahippocampal cortex (PHC). **(D)** Scene pair pattern similarity did not significantly predict saccade error in PPA (t(31) = −0.14, p = 0.890, d_z_ = 0.016, more distinct - less distinct = −0.28 degrees, 95% CI = [−4.36, 3.80]), RSC (t(31) = 0.90, p = 0.373, d_z_ = 0.098, more distinct - less distinct = 1.63 degrees, 95% CI = [−2.05, 5.31]), or PHC (t(31) = −1.32, p = 0.198, d_z_ = 0.15, more distinct - less distinct = −2.55 degrees, 95% CI = [−6.51, 1.41]; N = 32 participants, two-tailed paired t-tests). For A-D, small dots represent individual participants. Large points represent the across-participant mean ± SEM. **(E)** Whole-brain searchlight clusters where lower scene pair pattern similarity (more distinct representations) was associated with lower search error on valid trials. Clusters are visualized for exploratory purposes at an uncorrected p value threshold of 0.01. Clusters were found in the right lateral occipital lobe (left) and the right hippocampus (right). ns = p > 0.05; ** p < 0.01

**Supplementary Figure 5.**
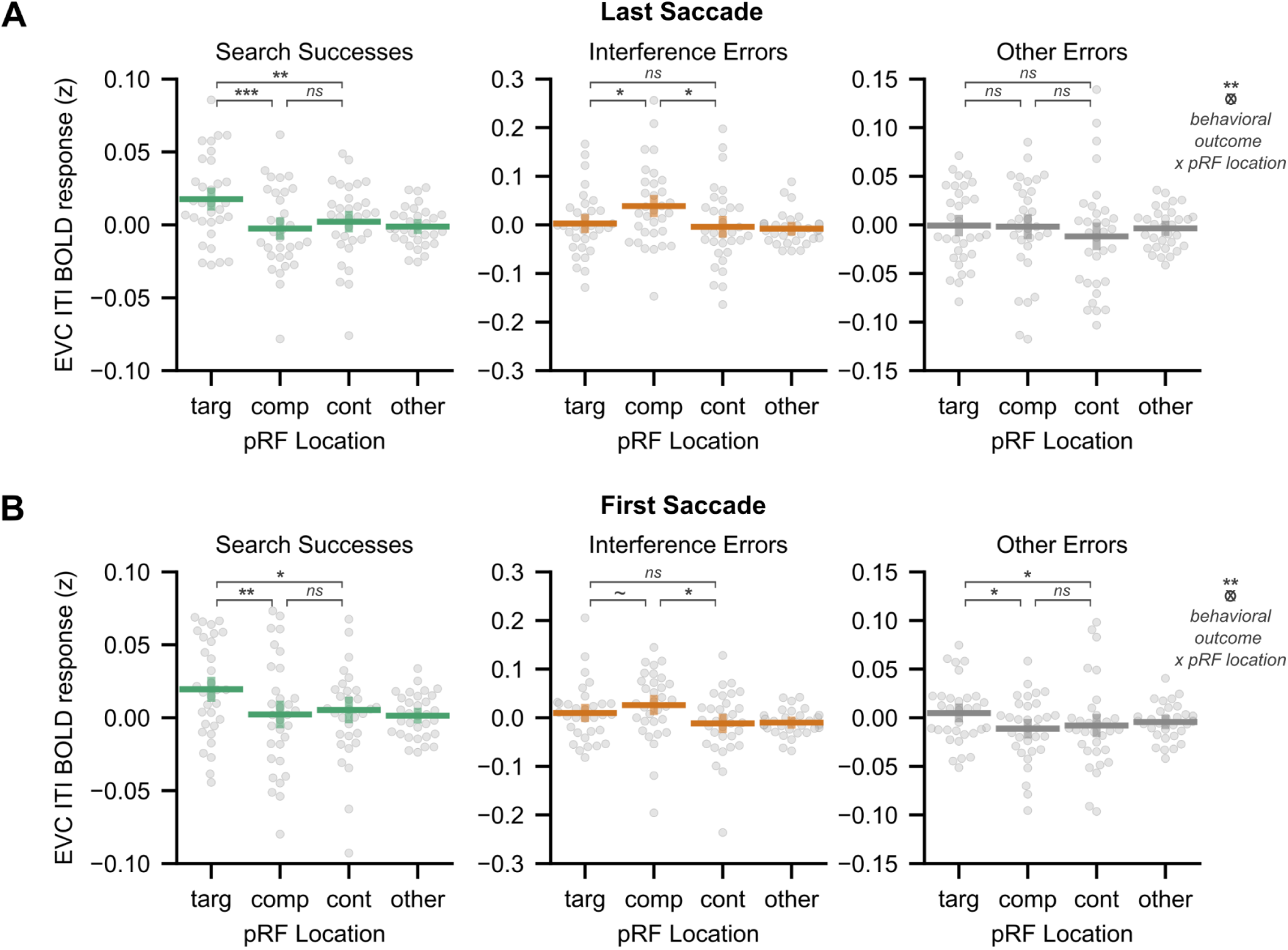
Preparatory coding in early visual cortex as a function of behavioral outcome defined by the last and first saccade of search. We directly compared early visual cortex (EVC) BOLD activity during the intertrial interval (ITI) at population receptive fields (pRFs) corresponding to the target location (targ), competitor location (comp), control location (cont), and all other locations (other) for three behavioral outcomes: search successes, interference errors, and other errors. **(A)** We defined search outcomes according to the participant’s last saccade of the search interval. Data from Figure 4D are replotted to visualize individual participants. **(B)** We defined search outcomes according to the participant’s first saccade of the search interval. Note that because outcomes are defined here with respect to the first saccade, trials may ultimately end in successful search even if the first saccade was an error. A linear mixed-effects model revealed an interaction between pRF location and behavioral outcome (F(6,75101) = 3.03, p = 0.006; N = 75144 trials across 32 participants; Type III F-test). This interaction was driven by preferential target activation for search successes, weak preferential competitor activation for interference errors (p = 0.112), and preferential target activation for other errors. Differences between this figure and panel A, notably in the other errors condition, are due to saccades that were initially errors but were ultimately corrected. Only pairwise statistical comparisons between target, competitor, and control locations are shown. For both panels, small dots represent individual participants. Horizontal lines represent the across-participant mean and vertical lines represent the SEM. ns = p > 0.05; *p < 0.05; **p < 0.01; *** p < 0.001

**Supplementary Figure 6.**
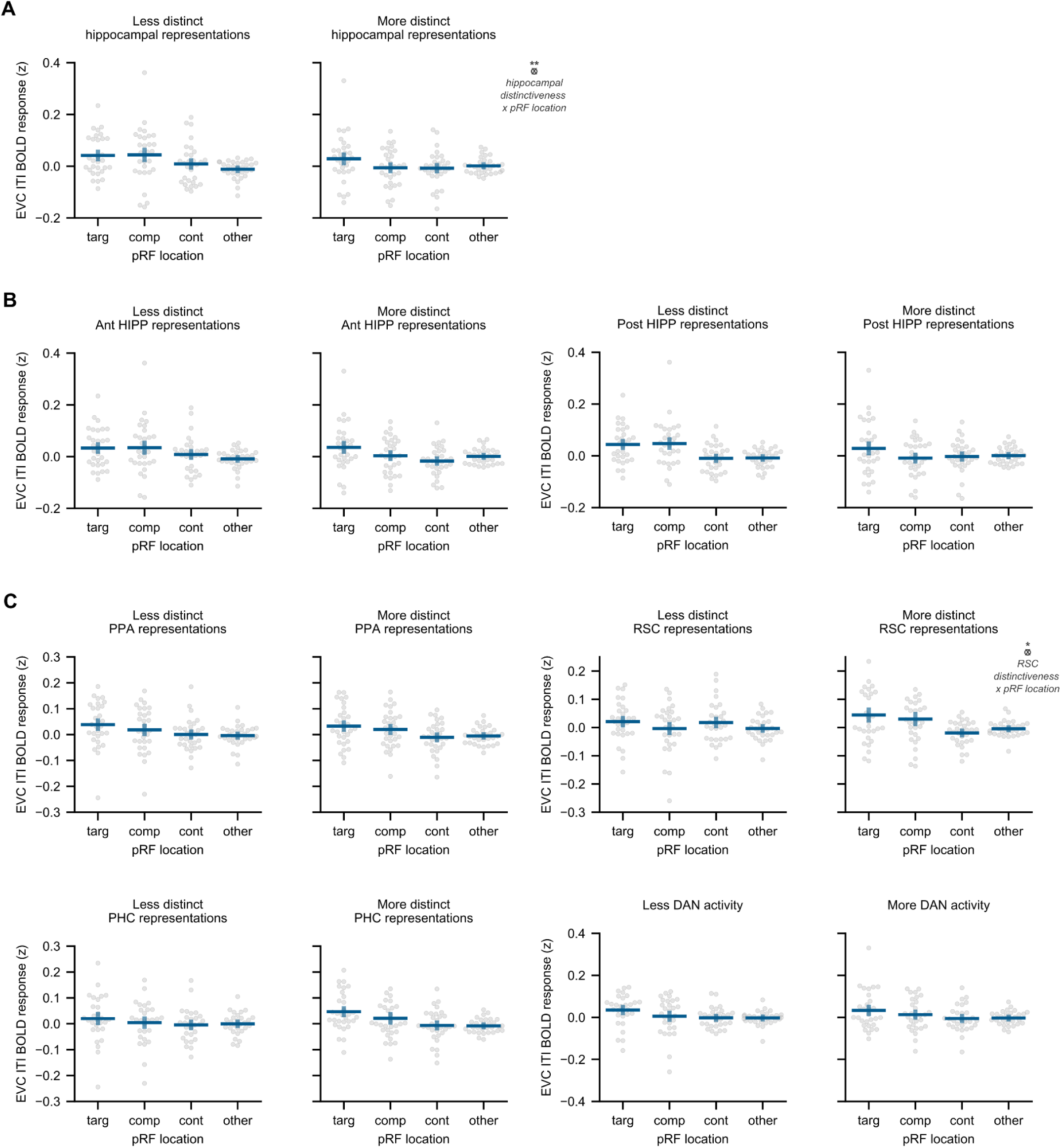
Brain regions outside the hippocampus are not associated with differences between target and competitor activation in visual cortex. **(A)** Data from Figure 5C is replotted to visualize individual participants. We repeated the analyses in Figure 5C in several brain regions outside the hippocampus. **(B)** We assessed scene distinctiveness in the anterior hippocampus (Ant HIPP) and posterior hippocampus (Post HIPP) separately. There was no significant interaction between hippocampal long-axis position, scene pair distinctiveness, and early visual cortex (EVC) population receptive field (pRF) location (F(3,39236) = 1.12, p = 0.340). We observed a qualitatively similar relationship in the anterior (F(3,19588.8) = 2.39, p = 0.066) and posterior hippocampus (F(3,19591.4) = 2.29, p = 0.077) as in the whole hippocampus: comparable target and competitor activation when scene representations were less distinct and higher target than competitor activation when scene representations were more distinct (Fig. 5C). **(C)** We assessed the effect of scene distinctiveness in parahippocampal place area (PPA), retrosplenial cortex (RSC), and parahippocampal cortex (PHC). For PPA and PHC, there was no significant interaction between EVC pRF location and PPA/PHC pattern distinctiveness (PPA: F(3,19617) = 0.44, p = 0.725; PHC: F(3,19587.5) = 1.40, p = 0.239). In RSC, there was an interaction (F(3,19587.7) = 3.17, p = 0.023), but the pattern of results was unlike those found in the hippocampus. Finally, we assessed the effect of univariate activity in the dorsal attention network (DAN). There was no significant interaction between EVC pRF location and univariate DAN activity (F(3,19592.3) = 1.155, p = 0.325). Note that we focus on the interaction with scene distinctiveness or univariate activity because main effects of target/competitor/control/other locations are not of primary interest. All statistics are Type III F-test run on N = 19656 trials across 32 participants. For all panels, small dots represent individual participants. Horizontal lines represent the across-participant mean and vertical lines represent the SEM. *p < 0.05.

## References

1. Carrasco, M. Visual attention: The past 25 years. Vision Res. 51, 1484–1525 (2011).

2. van Moorselaar, D. & Slagter, H. A. Inhibition in selective attention. Ann. N. Y. Acad. Sci. 1464, 204–221 (2020).

3. McAdams, C. J. & Maunsell, J. H. Effects of attention on orientation-tuning functions of single neurons in macaque cortical area V4. J. Neurosci. 19, 431–441 (1999).

4. Somers, D. C., Dale, A. M., Seiffert, A. E. & Tootell, R. B. H. Functional MRI reveals spatially specific attentional modulation in human primary visual cortex. Proc. Natl. Acad. Sci. 96, 1663–1668 (1999).

5. Liu, T., Pestilli, F. & Carrasco, M. Transient Attention Enhances Perceptual Performance and fMRI Response in Human Visual Cortex. Neuron 45, 469–477 (2005).

6. Sprague, T. C. & Serences, J. T. Attention modulates spatial priority maps in the human occipital, parietal and frontal cortices. Nat. Neurosci. 16, 1879–1887 (2013).

7. Desimone, R. & Duncan, J. Neural Mechanisms of Selective Visual Attention. Annu. Rev. Neurosci. 18, 193–222 (1995).

8. Kastner, S. & Ungerleider, L. G. Mechanisms of visual attention in the human cortex. Annu. Rev. Neurosci. 23, 315–341 (2000).

9. Norman, K. A., Newman, E., Detre, G. & Polyn, S. How inhibitory oscillations can train neural networks and punish competitors. Neural Comput. 18, 1577–1610 (2006).

10. Ritvo, V. J. H., Nguyen, A., Turk-Browne, N. B. & Norman, K. A. Differentiation and Integration of Competing Memories: A Neural Network Model. eLife 12, (2024).

11. Hulbert, J. C. & Norman, K. A. Neural differentiation tracks improved recall of competing memories following interleaved study and retrieval practice. Cereb. Cortex 25, 3994–4008 (2015).

12. Schlichting, M. L., Mumford, J. A. & Preston, A. R. Learning-related representational changes reveal dissociable integration and separation signatures in the hippocampus and prefrontal cortex. Nat. Commun. 6, 8151 (2015).

13. Favila, S. E., Chanales, A. J. H. & Kuhl, B. A. Experience-dependent hippocampal pattern differentiation prevents interference during subsequent learning. Nat. Commun. 6, 11066 (2016).

14. Chanales, A. J. H., Oza, A., Favila, S. E. & Kuhl, B. A. Overlap among spatial memories triggers divergence of hippocampal representations. Curr. Biol. 27, 2307–2317.e5 (2017).

15. Molitor, R. J., Sherrill, K. R., Morton, N. W., Miller, A. A. & Preston, A. R. Memory reactivation during learning simultaneously promotes dentate gyrus/CA2,3 pattern differentiation and CA1 memory integration. J. Neurosci. 41, 726–738 (2020).

16. Wanjia, G., Favila, S. E., Kim, G., Molitor, R. J. & Kuhl, B. A. Abrupt hippocampal remapping signals resolution of memory interference. Nat. Commun. 12, 4816 (2021).

17. Chun, M. M. & Phelps, E. A. Memory deficits for implicit contextual information in amnesic subjects with hippocampal damage. Nat. Neurosci. 2, 844–847 (1999).

18. Hannula, D. E. & Ranganath, C. The Eyes Have It: Hippocampal Activity Predicts Expression of Memory in Eye Movements. Neuron 63, 592–599 (2009).

19. Summerfield, J. J., Lepsien, J., Gitelman, D. R., Mesulam, M. M. & Nobre, A. C. Orienting attention based on long-term memory experience. Neuron 49, 905–916 (2006).

20. Aly, M. & Turk-Browne, N. B. Attention Stabilizes Representations in the Human Hippocampus. Cereb. Cortex 26, 783–796 (2016).

21. Córdova, N. I., Turk-Browne, N. B. & Aly, M. Focusing on what matters: Modulation of the human hippocampus by relational attention. Hippocampus 29, 1025–1037 (2019).

22. Ruiz, N. A., Meager, M. R., Agarwal, S. & Aly, M. The Medial Temporal Lobe Is Critical for Spatial Relational Perception. J. Cogn. Neurosci. 32, 1–16 (2020).

23. Rolls, E. T. Spatial view cells and the representation of place in the primate hippocampus. Hippocampus 9, 467–480 (1999).

24. Killian, N. J., Jutras, M. J. & Buffalo, E. a. A map of visual space in the primate entorhinal cortex. Nature 491, 761–764 (2012).

25. Mao, D. et al. Spatial modulation of hippocampal activity in freely moving macaques. Neuron 109, 3521–3534.e6 (2021).

26. Stokes, M., Atherton, K., Patai, E. Z. & Nobre, A. C. Long-term memory prepares neural activity for perception. Proc. Natl. Acad. Sci. 109, E360–E367 (2012).

27. Günseli, E. & Aly, M. Preparation for upcoming attentional states in the hippocampus and medial prefrontal cortex. eLife 9, e53191 (2020).

28. Ryan, J. D., Shen, K. & Liu, Z.-X. The intersection between the oculomotor and hippocampal memory systems: empirical developments and clinical implications. Ann. N. Y. Acad. Sci. 1464, 115–141 (2020).

29. Poskanzer, C. & Aly, M. Switching between External and Internal Attention in Hippocampal Networks. J. Neurosci. 43, 6538–6552 (2023).

30. Hutchinson, J. B. & Turk-Browne, N. B. Memory-guided attention: Control from multiple memory systems. Trends Cogn. Sci. 16, 576–579 (2012).

31. Aly, M. & Turk-Browne, N. B. How Hippocampal Memory Shapes, and Is Shaped by, Attention. in The Hippocampus from Cells to Systems (eds Hannula, D. E. & Duff, M. C.) 369–403 (Springer International Publishing, Cham, 2017). doi:10.1007/978-3-319-50406-3_12.

32. Posner, M. I. Orienting of attention. Q. J. Exp. Psychol. 32, 3–25 (1980).

33. Anderson, M. C., Green, C. & McCulloch, K. C. Similarity and inhibition in long-term memory: Evidence for a two-factor theory. J. Exp. Psychol. Learn. Mem. Cogn. 26, 1141–1159 (2000).

34. Chanales, A. J. H., Tremblay-McGaw, A. G., Drascher, M. L. & Kuhl, B. A. Adaptive Repulsion of Long-Term Memory Representations Is Triggered by Event Similarity. Psychol. Sci. 32, 705–720 (2021).

35. O’Reilly, R. C. & McClelland, J. L. Hippocampal conjunctive encoding, storage, and recall: Avoiding a trade-off. Hippocampus 4, 661–682 (1994).

36. Shapiro, M. L. & Olton, D. S. Hippocampal function and interference. in Memory systems 1994 87–117 (The MIT Press, Cambridge, MA, US, 1994).

37. Gandhi, S. P., Heeger, D. J. & Boynton, G. M. Spatial attention affects brain activity in human primary visual cortex. Proc. Natl. Acad. Sci. 96, 3314–3319 (1999).

38. Poppenk, J., Evensmoen, H. R., Moscovitch, M. & Nadel, L. Long-axis specialization of the human hippocampus. Trends Cogn. Sci. 17, 230–240 (2013).

39. Kosslyn, S. M., Thompson, W. L., Kim, I. J. & Alpert, N. M. Topographical representations of mental images in primary visual cortex. Nature 378, 496–498 (1995).

40. Kastner, S., Pinsk, M. A., De Weerd, P., Desimone, R. & Ungerleider, L. G. Increased activity in human visual cortex during directed attention in the absence of visual stimulation. Neuron 22, 751–61 (1999).

41. Breedlove, J. L., St-Yves, G., Olman, C. A. & Naselaris, T. Generative Feedback Explains Distinct Brain Activity Codes for Seen and Mental Images. Curr. Biol. 30, 2211–2224.e6 (2020).

42. Favila, S. E., Kuhl, B. A. & Winawer, J. Perception and memory have distinct spatial tuning properties in human visual cortex. Nat. Commun. 13, 5864 (2022).

43. Ritvo, V. J. H., Turk-Browne, N. B. & Norman, K. A. Nonmonotonic Plasticity: How Memory Retrieval Drives Learning. Trends Cogn. Sci. 23, 726–742 (2019).

44. Wammes, J., Norman, K. A. & Turk-Browne, N. Increasing stimulus similarity drives nonmonotonic representational change in hippocampus. eLife 11, e68344 (2022).

45. Chun, M. M. & Jiang, Y. Contextual cueing: implicit learning and memory of visual context guides spatial attention. Cognit. Psychol. 36, 28–71 (1998).

46. Fan, J. E. & Turk-Browne, N. B. Incidental Biasing of Attention From Visual Long-Term Memory. J. Exp. Psychol. Learn. Mem. Cogn. 42, 970–977 (2016).

47. Chun, M. M. & Turk-Browne, N. B. Interactions between attention and memory. Curr. Opin. Neurobiol. 17, 177–184 (2007).

48. Nobre, A. C. & Stokes, M. G. Premembering Experience: A Hierarchy of Time-Scales for Proactive Attention. Neuron 104, 132–146 (2019).

49. Nickel, A. E., Hopkins, L. S., Minor, G. N. & Hannula, D. E. Attention capture by episodic long-term memory. Cognition 201, 104312 (2020).

50. Wynn, J. S., Ryan, J. D. & Buchsbaum, B. R. Eye movements support behavioral pattern completion. Proc. Natl. Acad. Sci. 117, 6246–6254 (2020).

51. Hirschstein, Z. & Aly, M. Long-term memory and working memory compete and cooperate to guide attention. Atten. Percept. Psychophys. 85, 1517–1549 (2023).

52. Ryan, J. D., Althoff, R. R., Whitlow, S. & Cohen, N. J. Amnesia is a Deficit in Relational Memory. Psychol. Sci. 11, 454–461 (2000).

53. Goldfarb, E. V., Chun, M. M. & Phelps, E. A. Memory-Guided Attention: Independent Contributions of the Hippocampus and Striatum. Neuron 89, 317–324 (2016).

54. Kerrén, C., van Bree, S., Griffiths, B. J. & Wimber, M. Phase separation of competing memories along the human hippocampal theta rhythm. eLife 11, e80633 (2022).

55. Fernandez, C., Jiang, J., Wang, S.-F., Choi, H. L. & Wagner, A. D. Representational integration and differentiation in the human hippocampus following goal-directed navigation. eLife 12, e80281 (2023).

56. Jiang, J., Wang, S.-F., Guo, W., Fernandez, C. & Wagner, A. D. Prefrontal reinstatement of contextual task demand is predicted by separable hippocampal patterns. Nat. Commun. 11, 2053 (2020).

57. Ballard, I. C., Wagner, A. D. & Mcclure, S. M. Hippocampal pattern separation supports reinforcement learning. Nat. Commun. 10, 1073 (2019).

58. Yassa, M. A. & Stark, C. E. L. Pattern separation in the hippocampus. Trends Neurosci. 34, 515–525 (2011).

59. Dandolo, L. C. & Schwabe, L. Time-dependent memory transformation along the hippocampal anterior–posterior axis. Nat. Commun. 2018 91 9, 1205 (2018).

60. Cowan, E. T. et al. Time-dependent transformations of memory representations differ along the long axis of the hippocampus. Learn. Mem. 28, 329–340 (2021).

61. Zeidman, P. & Maguire, E. A. Anterior hippocampus: the anatomy of perception, imagination and episodic memory. Nat. Rev. Neurosci. 17, 173–182 (2016).

62. Stokes, M., Thompson, R., Cusack, R. & Duncan, J. Top-Down Activation of Shape-Specific Population Codes in Visual Cortex during Mental Imagery. J. Neurosci. 29, 1565–1572 (2009).

63. Bosch, S. E., Jehee, J. F. M., Fernandez, G. & Doeller, C. F. Reinstatement of Associative Memories in Early Visual Cortex Is Signaled by the Hippocampus. J. Neurosci. 34, 7493–7500 (2014).

64. Kok, P., Failing, M. F. & de Lange, F. P. Prior Expectations Evoke Stimulus Templates in the Primary Visual Cortex. J. Cogn. Neurosci. 26, 1546–1554 (2014).

65. Naselaris, T., Olman, C. A., Stansbury, D. E., Ugurbil, K. & Gallant, J. L. A voxel-wise encoding model for early visual areas decodes mental images of remembered scenes. NeuroImage 105, 215–228 (2015).

66. Woodry, R., Curtis, C. E. & Winawer, J. Feedback Scales the Spatial Tuning of Cortical Responses during Both Visual Working Memory and Long-Term Memory. J. Neurosci. 45, (2025).

67. Kuhl, B. A., Rissman, J., Chun, M. M. & Wagner, A. D. Fidelity of neural reactivation reveals competition between memories. Proc. Natl. Acad. Sci. 108, 5903–5908 (2011).

68. Lewis-Peacock, J. A. & Norman, K. A. Competition between items in working memory leads to forgetting. Nat. Commun. 5, 1–10 (2014).

69. Yonelinas, A. P. The hippocampus supports high-resolution binding in the service of perception, working memory and long-term memory. Behav. Brain Res. 254, 34–44 (2013).

70. Kolarik, B. S., Baer, T., Shahlaie, K., Yonelinas, A. P. & Ekstrom, A. D. Close but no cigar: Spatial precision deficits following medial temporal lobe lesions provide novel insight into theoretical models of navigation and memory. Hippocampus 28, 31–41 (2018).

71. Yu, Q., Teng, C. & Postle, B. R. Different states of priority recruit different neural representations in visual working memory. PLOS Biol. 18, e3000769 (2020).

72. Kok, P., Jehee, J. F. M. & de Lange, F. P. Less Is More: Expectation Sharpens Representations in the Primary Visual Cortex. Neuron 75, 265–270 (2012).

73. Hindy, N. C., Ng, F. Y. & Turk-Browne, N. B. Linking pattern completion in the hippocampus to predictive coding in visual cortex. Nat. Neurosci. 19, 665–667 (2016).

74. Kok, P. & Turk-Browne, N. B. Associative prediction of visual shape in the hippocampus. J. Neurosci. 38, 0163–18 (2018).

75. Tarder-Stoll, H., Baldassano, C. & Aly, M. The brain hierarchically represents the past and future during multistep anticipation. Nat. Commun. 15, 9094 (2024).

76. Finnie, P. S. B., Komorowski, R. W. & Bear, M. F. The spatiotemporal organization of experience dictates hippocampal involvement in primary visual cortical plasticity. Curr. Biol. 31, 3996–4008.e6 (2021).

77. Amer, T. & Davachi, L. Extra-hippocampal contributions to pattern separation. eLife 12, e82250 (2023).

78. Corbetta, M. & Shulman, G. L. Control of Goal-Directed and Stimulus-Driven Attention in the Brain. Nat. Rev. Neurosci. 3, 215–229 (2002).

79. Rosen, M. L., Stern, C. E., Michalka, S. W., Devaney, K. J. & Somers, D. C. Influences of Long-Term Memory-Guided Attention and Stimulus-Guided Attention on Visuospatial Representations within Human Intraparietal Sulcus. J. Neurosci. 35, 11358–11363 (2015).

80. Rosen, M. L., Stern, C. E., Michalka, S. W., Devaney, K. J. & Somers, D. C. Cognitive Control Network Contributions to Memory-Guided Visual Attention. Cereb. Cortex 26, 2059–2073 (2016).

81. Wimber, M., Alink, A., Charest, I., Kriegeskorte, N. & Anderson, M. C. Retrieval induces adaptive forgetting of competing memories via cortical pattern suppression. Nat. Neurosci. 18, 582–589 (2015).

82. Anderson, M. C. & Hulbert, J. C. Active Forgetting: Adaptation of Memory by Prefrontal Control. Annu. Rev. Psychol. 72, 1–36 (2021).

83. Kuhl, B. A., Dudukovic, N. M., Kahn, I. & Wagner, A. D. Decreased demands on cognitive control reveal the neural processing benefits of forgetting. Nat. Neurosci. 10, 908–914 (2007).

84. Yamins, D. L. K. et al. Performance-optimized hierarchical models predict neural responses in higher visual cortex. Proc. Natl. Acad. Sci. 111, 8619–8624 (2014).

85. Eickenberg, M., Gramfort, A., Varoquaux, G. & Thirion, B. Seeing it all: Convolutional network layers map the function of the human visual system. NeuroImage 152, 184–194 (2017).

86. Lindsay, G. W. Convolutional Neural Networks as a Model of the Visual System: Past, Present, and Future. J. Cogn. Neurosci. 33, 2017–2031 (2020).

87. Benson, N. C. & Winawer, J. Bayesian analysis of retinotopic maps. eLife 7, 0–45 (2018).

88. Himmelberg, M. M. et al. Cross-dataset reproducibility of human retinotopic maps. NeuroImage 244, 118609 (2021).

89. Li, H.-H., Sprague, T. C., Yoo, A. H., Ma, W. J. & Curtis, C. E. Joint representation of working memory and uncertainty in human cortex. Neuron 109, P3699–3712.e6 (2021).

90. Esteban, O. et al. Crowdsourced MRI quality metrics and expert quality annotations for training of humans and machines. Sci. Data 6, 30 (2019).

91. Esteban, O. et al. fMRIPrep: a robust preprocessing pipeline for functional MRI. Nat. Methods 16, 111–116 (2019).

92. Gorgolewski, K. et al. Nipype: A Flexible, Lightweight and Extensible Neuroimaging Data Processing Framework in Python. Front. Neuroinformatics 5, (2011).

93. Tustison, N. J. et al. N4ITK: Improved N3 Bias Correction. IEEE Trans. Med. Imaging 29, 1310–1320 (2010).

94. Avants, B. B., Epstein, C. L., Grossman, M. & Gee, J. C. Symmetric diffeomorphic image registration with cross-correlation: Evaluating automated labeling of elderly and neurodegenerative brain. Med. Image Anal. 12, 26–41 (2008).

95. Zhang, Y., Brady, M. & Smith, S. Segmentation of brain MR images through a hidden Markov random field model and the expectation-maximization algorithm. IEEE Trans. Med. Imaging 20, 45–57 (2001).

96. Dale, A. M., Fischl, B. & Sereno, M. I. Cortical surface-based analysis. I. Segmentation and surface reconstruction. NeuroImage 9, 179–194 (1999).

97. Klein, A. et al. Mindboggling morphometry of human brains. PLOS Comput. Biol. 13, e1005350 (2017).

98. Fonov, V., Evans, A., McKinstry, R., Almli, C. & Collins, D. Unbiased nonlinear average age-appropriate brain templates from birth to adulthood. NeuroImage 47, S102 (2009).

99. Jenkinson, M., Bannister, P., Brady, M. & Smith, S. Improved Optimization for the Robust and Accurate Linear Registration and Motion Correction of Brain Images. NeuroImage 17, 825–841 (2002).

100. Cox, R. W. & Hyde, J. S. Software tools for analysis and visualization of fMRI data. NMR Biomed. 10, 171–178 (1997).

101. Greve, D. N. & Fischl, B. Accurate and robust brain image alignment using boundary-based registration. NeuroImage 48, 63–72 (2009).

102. Power, J. D. et al. Methods to detect, characterize, and remove motion artifact in resting state fMRI. NeuroImage 84, 320–341 (2014).

103. Behzadi, Y., Restom, K., Liau, J. & Liu, T. T. A component based noise correction method (CompCor) for BOLD and perfusion based fMRI. NeuroImage 37, 90–101 (2007).

104. Satterthwaite, T. D. et al. An improved framework for confound regression and filtering for control of motion artifact in the preprocessing of resting-state functional connectivity data. NeuroImage 64, 240–256 (2013).

105. Lanczos, C. A Precision Approximation of the Gamma Function. J. Soc. Ind. Appl. Math. Ser. B Numer. Anal. 1, 86–96 (1964).

106. Abraham, A. et al. Machine learning for neuroimaging with scikit-learn. Front. Neuroinformatics 8, (2014).

107. Dumoulin, S. O. & Wandell, B. A. Population receptive field estimates in human visual cortex. NeuroImage 39, 647–660 (2008).

108. Dougherty, R. F. et al. Visual field representations and locations of visual areas v1/2/3 in human visual cortex. J. Vis. 3, 586–598 (2003).

109. Wandell, B., Dumoulin, S. & Brewer, A. a. Visual Field Maps in Human Cortex. Neuron 56, 366–383 (2007).

110. Rosenke, M., van Hoof, R., van den Hurk, J., Grill-Spector, K. & Goebel, R. A Probabilistic Functional Atlas of Human Occipito-Temporal Visual Cortex. Cereb. Cortex 31, 603–619 (2021).

111. Aly, M. & Turk-Browne, N. B. Attention promotes episodic encoding by stabilizing hippocampal representations. Proc. Natl. Acad. Sci. 113, E420–E429 (2016).

112. Yeo, B. T. T. et al. The organization of the human cerebral cortex estimated by intrinsic functional connectivity. J. Neurophysiol. 106, 1125–1165 (2011).

113. Barr, D. J., Levy, R., Scheepers, C. & Tily, H. J. Random effects structure for confirmatory hypothesis testing: Keep it maximal. J. Mem. Lang. 68, 255–278 (2013).

114. Bates, D., Kliegl, R., Vasishth, S. & Baayen, H. Parsimonious Mixed Models. Preprint at 10.48550/arXiv.1506.04967 (2018).

115. Luke, S. G. Evaluating significance in linear mixed-effects models in R. Behav. Res. Methods 49, 1494–1502 (2017).

116. Bates, D., Mächler, M., Bolker, B. & Walker, S. Fitting Linear Mixed-Effects Models Using lme4. J. Stat. Softw. 67, 1–48 (2015).

117. Kuznetsova, A., Brockhoff, P. B. & Christensen, R. H. B. lmerTest Package: Tests in Linear Mixed Effects Models. J. Stat. Softw. 82, 1–26 (2017).

118. Lenth, R. V. & Piaskowski, J. emmeans: Estimated Marginal Means, aka Least-Squares Means. (2026).

119. Favila, S.E. and Aly, M. Memory Guided Visual Search (MGVS). OpenNeuro. 10.18112/openneuro.ds008366.v1.0.0 (2026).

120. Favila, S. E. & Aly, M. Hippocampal mechanisms underlying the resolution of competition in memory and perception. Open Science Foundation. 10.17605/OSF.IO/C948A (2026).

